# β-adrenergic blockers increase cAMP and stimulate insulin secretion through a PKA/RYR2/TRPM5 pathway in pancreatic β-cells in vitro

**DOI:** 10.1101/2024.10.15.618403

**Authors:** Naoya Murao, Risa Morikawa, Yusuke Seino, Kenju Shimomura, Yuko Maejima, Yuichiro Yamada, Atsushi Suzuki

## Abstract

β-adrenergic blockers (β-blockers) are extensively used to inhibit β-adrenoceptor activation and subsequent cAMP production in many cell types. In this study, we characterized the effects of β-blockers on mouse pancreatic β-cells. Unexpectedly, high doses (100 μM) of β- blockers (propranolol and bisoprolol) led to a 5–10 fold increase in cAMP levels, enhanced intracellular influx, and stimulated a 2–4 fold increase in glucose-and glimepiride-induced insulin secretion in MIN6-K8 clonal β-cells and isolated mouse pancreatic islets. These effects were observed despite minimal expression of β-adrenoceptors in these cells. Mechanistically, cAMP increase led to ryanodine receptor 2 (RYR2) phosphorylation via protein kinase A (PKA), triggering Ca^2+^-induced Ca^2+^ release (CICR). CICR then activates transient receptor potential cation channel subfamily M member 5 (TRPM5), resulting in increased Ca^2+^ influx via voltage-dependent Ca^2+^ channels. These effects contradict the conventional understanding of the pharmacology of β-blockers, highlighting the variability in β-blocker actions depending on the experimental context.

**Graphical abstract:** 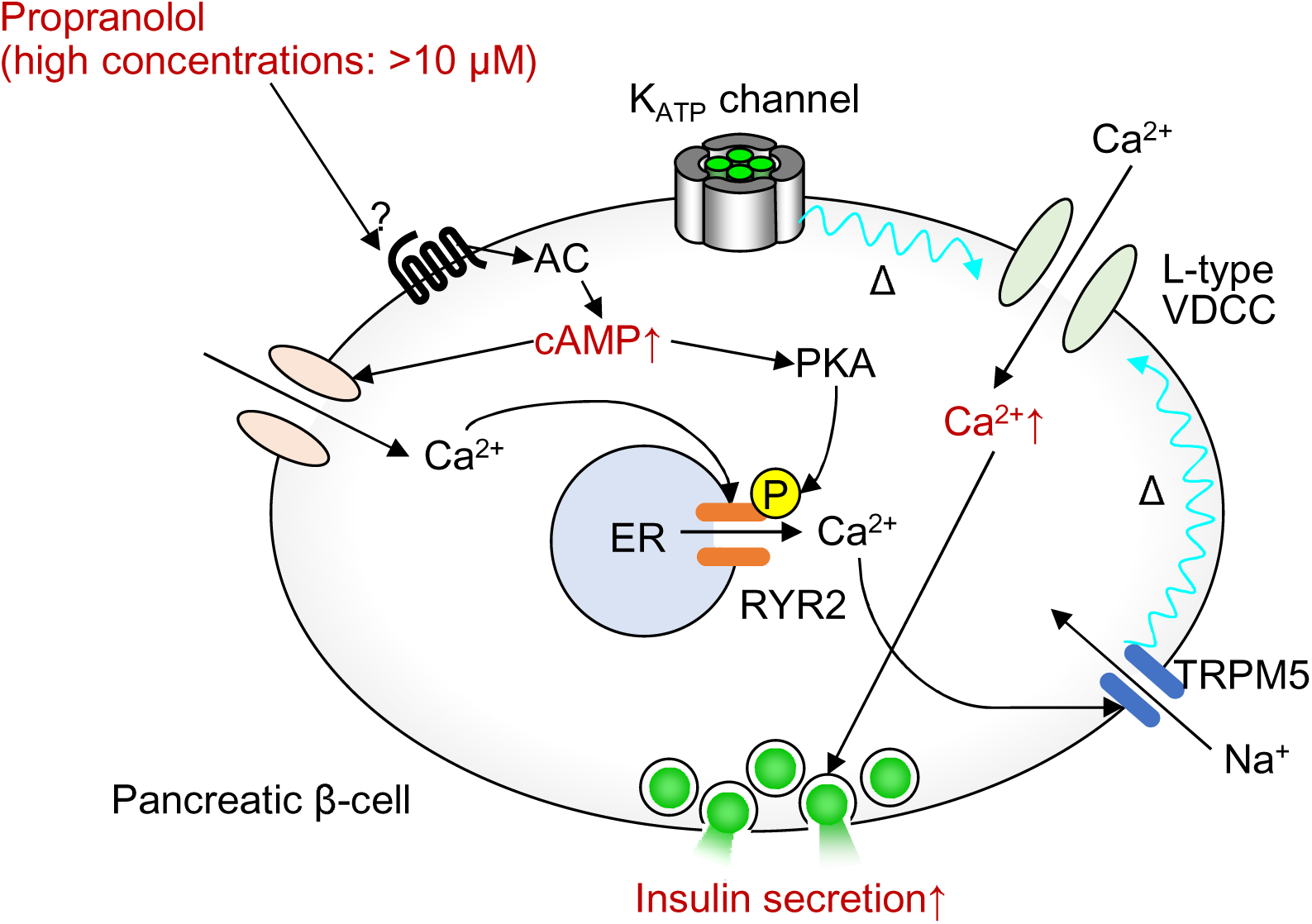

At high concentrations (> 10 μM), the β-adrenergic blocker propranolol paradoxically increased intracellular cAMP levels in pancreatic β-cells. This leads to PKA-induced RYR2 phosphorylation and extracellular Ca^2+^ influx, leading to CICR from the ER. CICR activated TRPM5, which augmented β-cell electrical activity, extracellular Ca^2+^ influx, and insulin secretion.

## Introduction

β-adrenergic blockers (β-blockers) are widely used for experimental and clinical purposes. The primary mechanism of β-adrenoceptors involves their coupling with Gαs proteins in trimeric G protein complexes, which subsequently stimulate adenylyl cyclase (AC) to generate adenosine 3’,5’-Cyclic Monophosphate (cAMP) (Brodde et al., 2006). β-blockers function by blocking the binding of catecholamines to β1-or β2-adrenoceptors, thus suppressing cAMP generation (Paul, 2021). Nevertheless, their mechanisms of action in cells outside the cardiovascular system are not well understood.

Pancreatic β-cells secrete insulin to maintain blood glucose levels within a physiological range. Glucose is the primary stimulator of insulin secretion; glucose metabolism leads to an increase in intracellular ATP levels, which inhibits ATP-sensitive K^+^ (K_ATP_) channels, resulting in membrane depolarization and opening of voltage-dependent Ca^2+^ channels, increasing intracellular Ca^2+^ ([Ca^2+^]_i_) levels and thereby stimulating insulin granule exocytosis (Henquin, 2000; Henquin, 2009).

Among the various adrenoceptor subtypes, β-cells primarily express α2A-adrenoceptors (Rorsman and Ashcroft, 2018). It is generally recognized that adrenaline suppresses insulin release from β-cells (Schuit and Pipeleers, 1986; Henquin, 2021; Sluga et al., 2022) by stimulating α2-adrenoceptors and the trimeric G protein Gαi/o, which in turn inhibits the production of adenosine 3’,5’-Cyclic Monophosphate (cAMP) by adenylyl cyclase (AC) (Straub and Sharp, 2012). Conversely, studies have shown that β-adrenoceptor agonists (β-agonists) increase insulin levels in the blood of mice and humans when administered in vivo. This effect is counteracted by β2-selective β-blockers (Porte, 1967; Cerasi et al., 1969; Imura et al., 1971; Loubatières et al., 1971; Ahren and Lundquist, 1981). However, these findings were not reproduced in vitro.

Here, we describe an unexpected effect of β-blockers on β-cells in vitro: when administered at high concentrations (> 10 μM), β-blockers significantly increased cAMP levels, [Ca^2+^]_i_, and insulin release in mouse β-cells. Our findings highlight the variable effects of β-blockers depending on the biological context, emphasizing the need for caution when using these compounds for experimental purposes.

## Materials and Methods

### Animals

C57BL/6JJcl (RRID:IMSR_JCL:JCL:MIN-0003) mice were purchased from CLEA Japan (Tokyo, Japan) and used for experiments at 19–20 weeks of age. NSY.B6-*Tyr*+, *A^y^*/Hos mice were a gift from Hoshino Laboratory Animals (Ibaraki, Japan) and were used for experiments at 19–20 weeks of age. Mice were maintained under specific-pathogen-free conditions at 23 ± 2 °C and 55 ± 10% relative humidity with 12-hour light-dark cycles (8am–8pm), with free access to water and standard chow CE-2 (CLEA Japan, Tokyo, Japan). The health status of the mice was checked regularly. All experiments were performed using male mice. All animal experiments were performed with the approval APU22080 by the Institutional Animal Care and Use Committee of Fujita Health University, complying with the Guidelines for Animal Experimentation at Fujita Health University and current Japanese guidelines and regulations for scientific and ethical animal experimentation (Kagiyama et al., 2006).

### Cell lines

MIN6-K8 cells were generated as previously described (Iwasaki et al., 2010) and were kindly provided by Professor Junichi Miyazaki (Osaka University). *Kcnj11*^-/-^ β-cells (clone name: *Kcnj11*^-/-^βCL1) were established by sub-cloning MIN6-K8 cells transfected with Cas9 nickase and guide RNA pairs targeting mouse *Kcnj11*, as described previously (Oduori et al., 2020). All cells were cultured in Dulbecco’s modified Eagle’s medium (DMEM) containing 4500 mg/L glucose (Sigma-Aldrich, St. Louis, MO, USA, Cat# D5796) supplemented with 10% fetal bovine serum (FBS) (BioWest, Nuaillé, France, Cat# S1400-500) and 5 ppm 2-mercaptoethanol. The cells were maintained at 37 °C with 5% CO_2_.

### Reagents

2.8 mM glucose-supplemented Krebs-Ringer bicarbonate buffer-HEPES (2.8G-KRBH) was prepared by supplementing KRBH (133.4 mM NaCl, 4.7 mM KCl, 1.2 mM KH_2_PO_4_, 1.2 mM MgSO_4_, 2.5 mM CaCl_2_, 5 mM NaHCO_3_, 10 mM HEPES, pH 7.4) with 0.1% bovine-serum albumin (Sigma-Aldrich, Cat# A6003) and 2.8 mM glucose. Ca^2+^-free KRBH was formulated by replacing CaCl_2_ with an equimolar MgCl_2_ and adding 0.2 mM EGTA (NACALAI TESQUE, Kyoto, Japan, Cat# 15214-21). For extracellular Na^+^ depletion experiments, control buffer (150 mM NaCl, 5 mM KCl, 1.2 mM KH_2_PO_4_, 1 mM MgCl_2_, 2.2 mM CaCl_2_, 10 mM HEPES, pH 7.4) and Na^+^-free buffer [150 mM N-Methyl-D-glucamine (Tokyo Chemical Industry, Tokyo, Japan, CAS: 6284-40-8, Cat# M0227), 5 mM KCl, 1.2 mM KH_2_PO_4_, 1 mM MgCl_2_, 2.2 mM CaCl_2_, 10 mM HEPES, pH 7.4] were prepared. 30 mM Na^+^ buffer was formulated by mixing the control buffer and Na^+^-free buffer at a ratio of 1:4 and supplementing 0.1% bovine-serum albumin (Sigma-Aldrich, Cat# A6003) and 2.8 mM glucose.

The following reagents were added to KRBH during the stimulation period: 11.1 mM glucose, (±)-propranolol hydrochloride (FUJIFILM Wako Pure Chemical, Osaka, Japan, CAS: 318-98-9, Cat# 168-28071), carteolol hydrochloride (Tokyo Chemical Industry, CAS: 51781-21-6, Cat# C3015), nadolol (Sigma-Aldrich, CAS: 42200-33-9, Cat# N1892), timolol maleate (Tokyo Chemical Industry, CAS: 26921-17-5, Cat# T2905), bisoprolol hemifumarate (Tokyo Chemical Industry, CAS: 104344-23-2, Cat# B3994), metoprolol tartrate (Tokyo Chemical Industry, Tokyo, Japan, CAS: 56392-17-7, Cat# M2555), glimepiride (Tokyo Chemical Industry, CAS: 93479-97-1, Cat# G0395), and carbamylcholine chloride (carbachol) (Tokyo Chemical Industry, CAS: 51-83-2, Cat# C0596).

The following reagents were added to KRBH during the pre-incubation and stimulation periods: nifedipine (FUJIFILM Wako Pure Chemical, Osaka, Japan, Cat# 14505781), thapsigargin (FUJIFILM Wako Pure Chemical, Osaka, Japan, CAS: 67526-95-8, Cat# 209-17281), MDL-12330A hydrochloride (Sigma-Aldrich, CAS: 40297-09-4, Cat# M184), SQ22536 (Selleck Chemicals, Houston, TX, USA, CAS: 17318-31-9, Cat# S8283), H-89 hydrochloride (Cayman Chemical, Ann Arbor, MI, USA, CAS: 130964-39-5, Cat# 10010556), phentolamine mesylate (Tokyo Chemical Industry, CAS: 65-28-1, Cat# P1985), atipamezole hydrochloride (FUJIFILM Wako Pure Chemical, Osaka, Japan, CAS: 104075-48-1, Cat# 015-25331), dantrolene sodium salt hydrate (Tokyo Chemical Industry, CAS: 14663-23-1, Cat# D3996), triphenylphosphine oxide (Tokyo Chemical Industry, CAS: 791-28-6, Cat# T0625), YM-254890 (FUJIFILM Wako Pure Chemical, Osaka, Japan, CAS: 568580-02-9, Cat# 257-00631), t-butylhydroquinone (Tokyo Chemical Industry, CAS: 1948-33-0, Cat# 027-07212), and xestospongin C (FUJIFILM Wako Pure Chemical, Osaka, Japan, CAS: 244-00721, Cat# C0598). The reagents used for stimulation were stored as a 1000× concentrate in dimethyl sulfoxide (DMSO) (FUJIFILM Wako Pure Chemical, Osaka, Japan, Cat# 041-29351) and diluted with KRBH shortly before the experiment. An equal volume of DMSO was added to the vehicle control.

### Isolation of pancreatic islets from mice

Digesting solution was formulated by supplementing 0.1w/v% Collagenase from Clostridium histolyticum (Sigma-Aldrich, Cat# C6885) to Hanks’ balanced salt solution (136.9 mM NaCl, 5.4 mM KCl, 0.8 mM MgSO_4_, 0.3 mM Na_2_HPO_4_, 0.4 mM KH_2_PO_4_, 4.2 mM NaHCO_3_, 10 mM HEPES, 1.3 mM CaCl_2_, 2 mM glucose). The mice were euthanized by isoflurane exposure. Pancreas was digested by 10-min incubation at 37 °C following intraductal injection of digesting solution. Islets were hand-picked and separated from exocrine tissues, transferred to 60-mm non-treated plates (AGC Techno Glass, Shizuoka, Japan, Cat# 1010-060), and cultured overnight in RPMI-1640 (Sigma-Aldrich, Cat# R8758) supplemented with 10% FBS (BioWest, Nuaillé, France, Cat# S1400-500) and 1% penicillin-streptomycin solution (FUJIFILM Wako Pure Chemical, Osaka, Japan, Cat# 168-23191) at 37 °C and 5% CO_2_ before the experiments.

### Insulin secretion and content

Insulin secretion was measured using the static incubation method, as described previously (Murao et al., 2022; Murao et al., 2024b). To measure insulin secretion from cell lines, the cells were seeded in 24-well plates (Corning, Glendale, AZ, USA, Cat# 353047) at a density of 5 × 10^5^ cells/well and cultured for 48 h. The cells were pre-incubated for 30 min with 300 μL/well of 2.8G-KRBH and subsequently stimulated with 300 μL/well of 2.8G-KRBH containing the specified stimulations for 30 min at 37 °C. The supernatant was collected for the measurement of secreted insulin, after which the cells were extracted with 0.1% Triton-X in 2.8G-KRBH for the measurement of insulin content. Insulin was quantified using a homogeneous time-resolved fluorescence assay (HTRF) Insulin Ultrasensitive kit (Revvity, Waltham, MA, USA, Cat# 62IN2PEH) in accordance with the manufacturer’s instructions. To measure insulin secretion from islets, overnight cultured islets were rinsed twice with 2.8G-KRBH, followed by pre-incubation with 2.8G-KRBH for 30 min at 37 °C. Size-matched islets were hand-picked and dispensed in a 96-well plate (Corning, Cat# 353072) at 5 islets/well. KRBH (100 μL/well) containing the specified stimulations was added and incubated for 30 min at 37 °C. The supernatant was subjected to insulin quantification as described above.

### Measurement of inositol 1-phosphate (IP1)

Intracellular inositol 1-phosphate content was measured as described previously (Murao et al., 2024a). Briefly, MIN6-K8 cells were seeded in a 96-well plate (Corning, Cat# 353072) at a density of 1.0 × 10^5^ cells/well and cultured for 48h. The measurement was performed using IP1 assay buffer (146 mM NaCl, 4.2 mM KCl, 50 mM LiCl, 1 mM CaCl2, 0.5 mM MgCl2, 10 mM HEPES, 0.1% BSA, pH7.4) supplemented with 2.8 mM glucose (2.8G-IP1 assay buffer). The cells were pre-incubated for 30 min with 30 μL/well of 2.8G-IP1 assay buffer, followed by the addition of 10 μL/well of IP1 assay buffer containing the specified stimulations at 4× concentration, and incubation for another 30 min at 37 °C. The reaction was terminated by the addition of 10 μL/well lysis buffer. The lysate (30 μL/well) was transferred to a 384-well plate (Corning, Cat #3826) and subjected to IP1 quantification using the HTRF IP-One Gq Detection Kit (Revvity, Waltham, MA, USA, Cat# 62IPAPEB) according to the manufacturer’s instructions.

### Measurement of cAMP

To prevent cAMP hydrolysis, 2.8G-KRBH was supplemented with 100 μM isobutylmethylxanthine (IBMX) (FUJIFILM Wako Pure Chemical, CAS: 28822-58-4, Cat# 095-03413) for pre-incubation and stimulation. To measure cAMP levels in MIN6-K8 cells, the cells were seeded in a 96-well plate (Corning, Cat# 353072) at a density of 1.0 × 10^5^ cells/well and cultured for 48h. The cells were pre-incubated for 30 min with 40 μL/well of 2.8G-KRBH, followed by the addition of 20 μL/well of 2.8G-KRBH containing the specified stimulations at 4× concentration, and incubation for another 30 min at 37 °C. The reaction was terminated by the addition of 20 μL/well lysis buffer. The lysate (20 μL/well) was transferred to a 384-well plate (Corning, Cat #3826) and subjected to cAMP quantification using the HTRF cAMP Gs Dynamic Detection Kit (Revvity, Waltham, MA, USA, Cat# 62AM4PEB) according to the manufacturer’s instructions.

To measure cAMP levels in islets, overnight cultured islets were rinsed twice with 2.8G-KRBH. Size-matched islets were hand-picked and dispensed in a 96-well plate (Corning, Cat# 353072) at 5 islets/well, followed by pre-incubation with 30 μL/well of 2.8G-KRBH for 30 min at 37 °C. Subsequently, 10 μL/well of 2.8G-KRBH containing the specified stimulations at 4× concentration was added to the wells and incubated for another 30 min at 37 °C. The reaction was terminated by the addition of 15 μL/well lysis buffer. The lysate (20 μL/well) was transferred to a 384-well plate and subjected to cAMP quantification, as described above.

### Ca^2+^ imaging

Ca^2+^ imaging was performed as previously described (Murao et al., 2024b). Briefly, MIN6-K8 cells were seeded in a 35 mm glass-bottom dish (Matsunami Glass, Osaka, Japan, Cat# D11530H) at a density of 1.28 × 10^5^ cells/dish and cultured for 48 h. Subsequently, the cells were loaded with 1 μM Fluo-4 AM (Dojindo, Kumamoto, Japan, Cat# F312) in 2.8G-KRBH for 20 min at 37 °C in room air. The cells were loaded with 1 mL of fresh 2.8G-KRBH, and basal recordings were performed for 300 s (from time-300 to 0). Following the addition of 1 mL KRBH supplemented with stimulations at a 2× concentration, recordings were resumed for another 600 s (from time 0 to 600). Time-lapse images were obtained using a Zeiss LSM 980 Airyscan2 inverted confocal laser scanning super-resolution microscope equipped with a Plan Apo 40×, 1.4 Oil DICII objective lens (Carl Zeiss Microscopy, Jena, Germany). The cells were excited at 488 nm laser with 0.3% output power, and fluorescence emission was measured at 508-579 nm. The obtained images were analyzed using the ZEN 3.0 imaging software (Carl Zeiss Microscopy, Jena, Germany, RRID:SCR_021725). Cells were randomly chosen for analysis for each stimulation, and the number of cells analyzed is indicated in the figure legends. The fluorescence intensity of the entire cell body (F) was monitored and normalized to the average fluorescence intensity between-300 and 0 s (F0).

### Gene silencing using small interfering RNA (siRNA)

Gene silencing was performed as previously described (Murao et al., 2024b). The following siRNAs were purchased from Dharmacon (Lafayette, CO, USA): *Adra2a* (Cat# M-062217-01-0005), *Itpr1* (Cat# M-040933-01-0005), *Trpm4* (Cat# M-056098-01-0005), *Trpm5* (Cat# M-046645-01-0005), and non-targeting siRNA (Cat# D-001206-14-50). Briefly, siRNAs were reverse-transfected using DharmaFECT 2 transfection reagent (Dharmacon, Lafayette, CO, USA, Cat# T-2002-03) according to the manufacturer’s instructions. The final concentrations of siRNA and DharmaFECT 2 were 40 nM and 0.4%, respectively. For insulin secretion experiments, cells were seeded in 24-well plates (Corning, Cat# 353047) at 5 × 10^5^ cells/500 μL/well. For immunoblotting, cells were seeded in 12-well plates (Corning, Cat# 353043) at 1 × 10^6^ cells/mL/well.

### Reverse transcription quantitative polymerase chain reaction (RT-qPCR)

RT-PCR was performed as previously described (Murao et al., 2024b). Briefly, cDNA was prepared using CellAmp Direct Lysis and RT set (Takara Bio, Shiga, Japan, Cat# 3737S/A) according to the manufacturer’s instructions. Quantitative real-time PCR was performed on a QuantStudio 7 Flex system (Thermo Fisher Scientific, Waltham, MA, USA, RRID:SCR_020245) using TaqMan Universal Master Mix II with UNG (Thermo Fisher Scientific, Waltham, MA, USA, Cat# 4440038) and Taqman probes as follows: *Adra2a* (Cat# Mm07295458_s1), *Itpr1* (Cat# Mm00439907_m1), *Trpm4* (Cat# Mm00613173_m1), *Trpm5* (Cat# Mm01129032_m1), and TATA-box binding protein (*Tbp*, Cat# Mm01277042_m1). Relative gene expression was calculated using the 2^-ΔΔCT^ method and normalized to *Tbp*.

### Immunoblotting

Immunoblotting was performed as previously described (Murao et al., 2024b), with slight modifications. Briefly, cells were lysed with 50 μL/well RIPA buffer supplemented with a complete protease inhibitor cocktail (Sigma-Aldrich, Cat# 11697498001) and PhosSTOP (Roche, Basel, Switzerland, Cat# 4906845001). For the detection of ADRA2A, the lysate was separated on a 7.5% polyacrylamide-SDS gel and transferred to a PVDF membrane. The membranes were probed with polyclonal anti-ADRA2A rabbit antibody (Thermo Fisher Scientific, Waltham, MA, USA, Cat# PA1-048, RRID:AB_2225243) overnight at 4 °C. To detect RYR2 phosphorylation, the lysate was separated on a 5% polyacrylamide-SDS gel and transferred to a PVDF membrane. The membranes were probed with either an anti-RYR2 rabbit monoclonal antibody (Abcam, Cambridge, UK, Cat# EPR26288-70, 1:300) or an anti-phospho RyR2 (Ser2808) rabbit polyclonal antibody (Millipore, Burlington, MA, USA, Cat# ABS2231, 1:300) overnight at 4 °C. The membrane was then probed with HRP-conjugated polyclonal swine anti-rabbit antibody (Agilent Technologies, Santa Clara, CA, USA, Cat# P0399, RRID:AB_2617141, 1:2000) and visualized using ECL Prime (Cytiva, Buckinghamshire, UK, Cat# RPN2232). To visualize the whole protein, the transferred membrane was stained with Ponceau-S solution (Beacle, Kyoto, Japan, Cat# BCL-PSS-01). Images were taken using ImageQuant 800 (Cytiva, Buckinghamshire, UK). The images were quantified using ImageJ (version 1.53k, https://imagej.nih.gov/ij/index.html, RRID:SCR_003070).

### Statistical Analysis

Sample sizes were estimated from the expected effect size based on previous experiments. No randomization or blinding was used. For cell line experiments, including insulin secretion, IP1 measurement, cAMP measurement, RT-qPCR, and immunoblotting, n represents the number of biological replicates of cells grown in different wells of the same multiwell plate. Results were confirmed by 2–3 independent experiments performed on different days using different cell passages. For islet experiments, n represents the number of wells, each containing five islets. The number of mice used is indicated in the figure legends. Results were confirmed by two independent experiments performed on different days using different mice. For Ca^2+^ imaging, n represents the number of different single cells analyzed. Data are shown as the mean ± standard deviation (SD) along with the plot of individual data points. For statistical comparisons between two groups, a two-tailed unpaired Welch’s unpaired t-test was used. For comparisons between more than three groups, Welch’s one-way analysis of variance (ANOVA) was followed by pairwise comparisons corrected using Dunnett’s method. Normality of the distribution was confirmed by the Shapiro-Wilk test. P-values are indicated in the figures. P-values less than 0.05 were considered statistically significant. The statistical analyses used are indicated in the figure legends. Statistical analyses were performed using GraphPad Prism 10 (GraphPad Software, Boston, MA, USA, RRID:SCR_002798).

## Results

### β-Blockers amplify insulin secretion from MIN6-K8 β-cell lines

To evaluate the impact of β-blockers on insulin release, we used MIN6-K8, a mouse β-cell line (Iwasaki et al., 2010). The cells were stimulated with propranolol (non-selective β- blocker) and bisoprolol (β1-selective blocker). In the presence of stimulatory levels (11.1 mM) of glucose, both propranolol and bisoprolol dose-dependently increased insulin secretion at concentrations greater than 10 μM and 100 μM, respectively (Figure 1, A–B).

**Figure 1.**
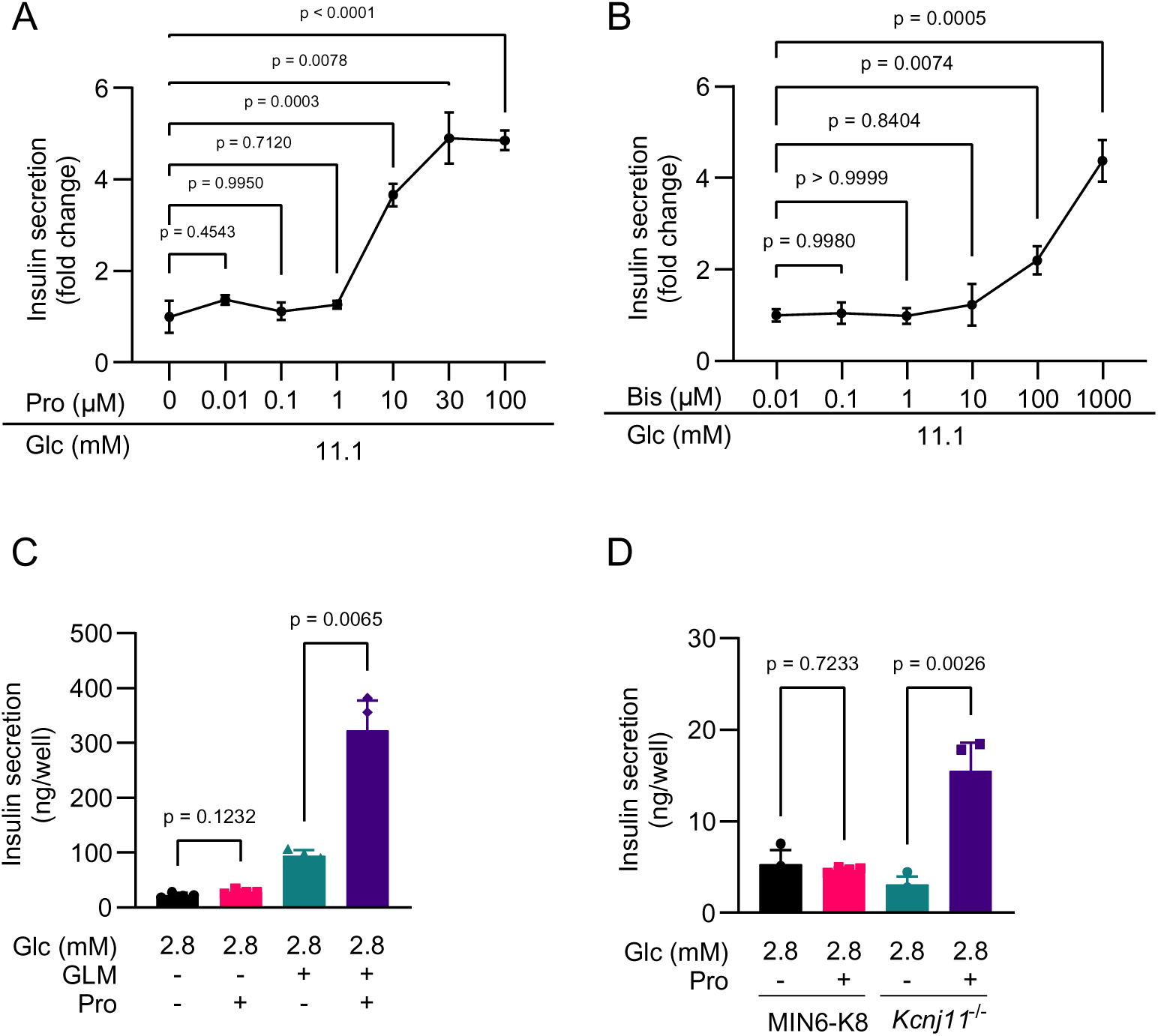
Effect of β-blockers on glucose-and glimepiride-induced insulin secretion in MIN6-K8 cells. A. Dose-dependent effect of propranolol on glucose-induced insulin secretion. n = 4. B. Dose-dependent effect of bisoprolol on glucose-induced insulin secretion. n = 4. C. Requirement of glimepiride for propranolol-stimulated insulin secretion at 2.8 mM glucose. n = 4. D. Propranolol-stimulated insulin secretion in a K_ATP_ channel-deficient β-cell line (*Kcnj11*^-/-^). n = 4. All experiments were conducted using MIN6-K8 cells unless otherwise specified. Data are presented as mean ± standard deviation (SD). Glc, glucose. The reagents were added to achieve the following final concentrations unless otherwise specified: glimepiride (GLM), 1 μM; and propranolol (Pro), 100 μM. Statistical comparisons were performed using Welch’s one-way ANOVA with Dunnett’s post-hoc test.

At basal glucose levels (2.8 mM), propranolol had no effect on insulin secretion (Figure 1C) or insulin content (Figure S1A). When co-administered with sulfonylurea (K_ATP_ channel inhibitor) glimepiride, propranolol amplified insulin secretion even at 2.8 mM glucose (Figure 1C). To evaluate whether propranolol-stimulated insulin secretion (propranolol-SIS) results from the promotion of K_ATP_ channel closure, its efficacy was examined in *Kcnj11*^-/-^ β- cells. As *Kcnj11* encodes the pore-forming subunit of the K_ATP_ channel, *Kcnj11*^-/-^ β-cells lack K_ATP_ channel activity, and their cell membrane is continuously depolarized regardless of extracellular glucose levels (Oduori et al., 2020). Propranolol increased insulin secretion in *Kcnj11*^-/-^ β-cells even at 2.8 mM glucose (Figure 1D), indicating that the primary mechanism for propranolol-stimulated insulin secretion is not through facilitating K_ATP_ channel closure. Rather, our results suggest that membrane depolarization by high glucose, sulfonylureas, or *Kcnj11* knockout is permissive for propranolol-SIS. Consequently, in subsequent experiments, β-blocker-stimulated insulin secretion (β-blocker-SIS) was examined in the presence of high glucose or glimepiride. We found that a wide range of β-blockers, both non-selective and β1- selective, enhanced insulin release, with nadolol as the sole exception (Figure S1B).

Therefore, the capacity to promote insulin secretion is likely a common feature among compounds in the β-blocker class.

### **β**-Blockers increase cAMP levels in MIN6-K8 **β**-cell lines

To elucidate the mechanism of β-blocker-SIS, we investigated whether β-blockers modulate canonical GPCR signaling pathways in MIN6-K8 cells.

Propranolol and bisoprolol dose-dependently increased intracellular cAMP levels in MIN6-K8 cells at concentrations that were effective for insulin secretion (Figure 2, A–B). Two adenylyl cyclase (AC) inhibitors, MDL-12330A and SQ-22536, suppressed cAMP accumulation induced by 100 μM propranolol and 1 mM bisoprolol (Figure 2C and S2A), indicating that β-blocker-induced cAMP accumulation is mediated by adenylyl cyclase activation. Furthermore, these inhibitors also abolished propranolol-SIS (Figure 2, D–E), indicating that cAMP production via AC is necessary for propranolol-SIS. Notably, propranolol-SIS was also abolished by H-89, a protein kinase A (PKA) inhibitor (Figure 2F), suggesting that the cAMP/PKA pathway is essential for propranolol-SIS.

**Figure 2.**
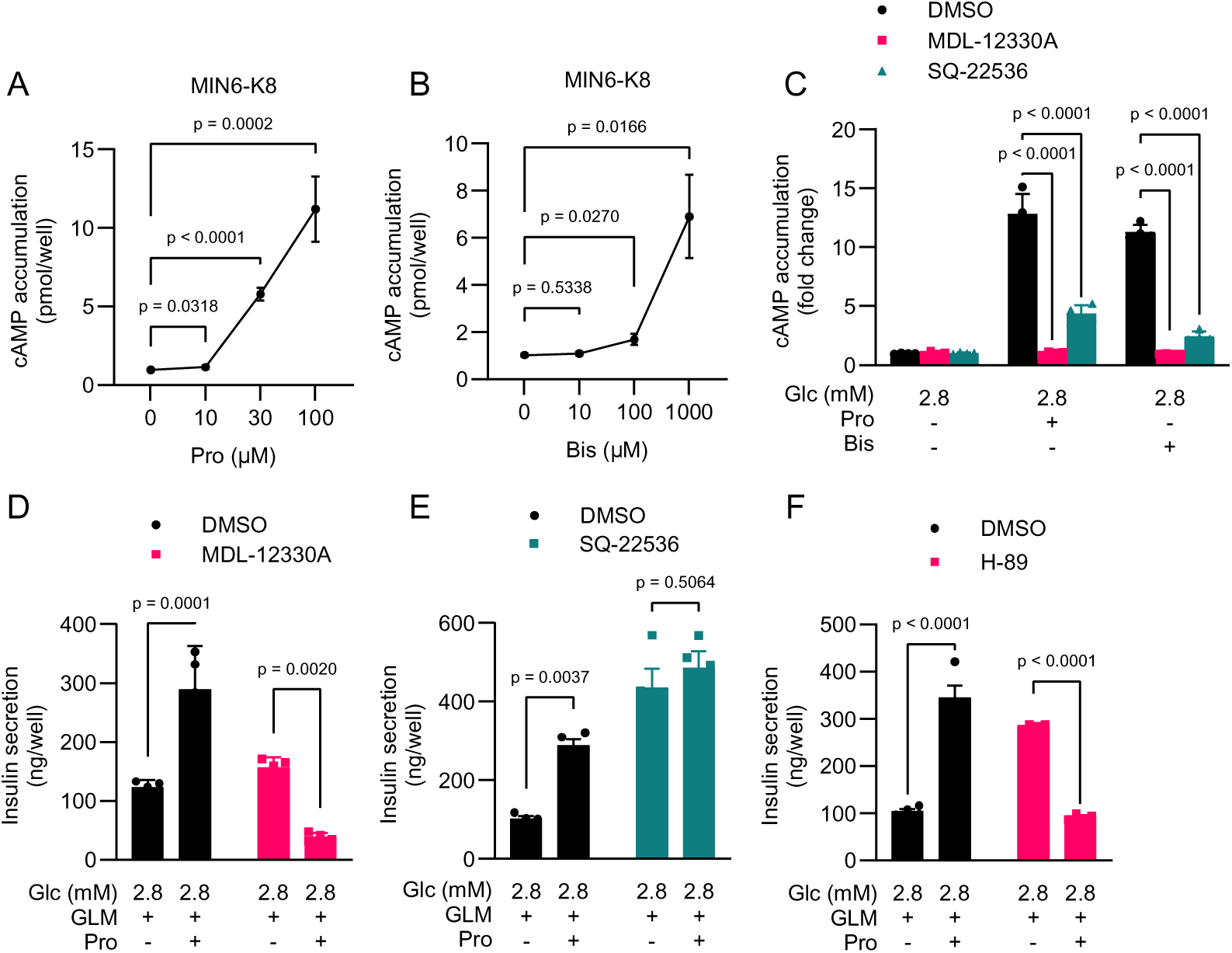
Characterization of β-blocker-stimulated cAMP elevation and insulin secretion in MIN6-K8 cells. A. Effect of propranolol on intracellular cAMP accumulation. n = 4. B. Effect of bisoprolol on intracellular cAMP accumulation. n = 4. C. Effects of AC inhibitors on β-blocker-stimulated cAMP elevation. n = 4. The data are presented as fold change over 2.8 mM glucose alone. See Figure S1B for the original values. D. –E. Effects of AC inhibitors on propranolol-SIS. MDL-12330A and SQ-22536 were used for D and E, respectively. n = 4. F. Effect of H-89 on propranolol-SIS. n = 4. All experiments were conducted using MIN6-K8 cells. Data are presented as mean ± SD. Glc, glucose. The reagents were added to achieve the following final concentrations unless otherwise specified: glimepiride (GLM) – 1 μM, propranolol (Pro) – 100 μM, bisoprolol (Bis) – 100 μM, MDL-12330A – 10 μM, SQ-22536 – 1 mM, and H-89 – 10 μM. Statistical comparisons were made using Welch’s one-way ANOVA with Dunnett’s post-hoc test for A and B, and two-way ANOVA with Šídák’s post-hoc test for C – F.

Subsequently, inositol 1-phosphate (IP1), a metabolite of inositol 1,4,5-trisphosphate (IP_3_), was measured to assess the activity of the Gαq/phospholipase Cβ (PLCβ) signaling pathway. IP1 levels remained unaltered by propranolol, although substantial accumulation of IP1 was confirmed using carbachol as a positive control (Figure S2B), indicating that propranolol does not activate the Gαq/PLCβ pathway. This finding is corroborated by the observation that YM-254890, a specific inhibitor of Gαq/11/14 (Takasaki et al., 2004; Patt et al., 2021), suppressed glimepiride-induced insulin secretion (Figure S2C), but minimally affected propranolol-SIS when expressed as a fold-change (Figure S2D). Collectively, these results demonstrate that propranolol-SIS involves the cAMP/PKA pathway, but not the Gαq/PLCβ pathway.

### Effect of **β**-blockers in mouse islets

We investigated whether these findings could be applied to primary β-cells. We observed enhancement of insulin release by propranolol in C57BL/6J (B6) mouse islets when exposed to 11.1 mM glucose (Figure 3, A–B) or glimepiride (Figure 3C). These findings are consistent with the results obtained using MIN6-K8 cells. NSY.B6-*Tyr*+, *A^y^* (NSY.B6*-A^y^*) is a spontaneous type 2 diabetes mouse strain established by crossing the NSY and B6J- *A^y^* strains, introducing the obesity-related agouti-yellow (*A^y^*) mutation in the agouti (a) gene into a diabetes-prone NSY background (Ohno et al., 2022). *A^y^* heterozygous male mice (NSY.B6*- A^y^*/*a*) exhibited obesity and hyperglycemia; wild-type male mice (NSY.B6*-a*/*a*) served as lean controls (Ohno et al., 2022; Murao et al., 2024b). At 2.8 mM glucose, no propranolol-SIS was observed in either strain (Figure 3, D–E), confirming the glucose dependency of propranolol-SIS (Figure 1G). Propranolol substantially increased insulin secretion in diabetic *A^y^*/*a* islets (Figure 3E), although it did not reach statistical significance in lean *a*/*a* islets (Figure 3D). Notably, propranolol also increased intra-islet cAMP levels in both strains (Figure 3, F–G). These findings indicate that the observed effects are likely a general characteristic of β-blockers on β-cells rather than an artifact specific to a cell line.

**Figure 3.**
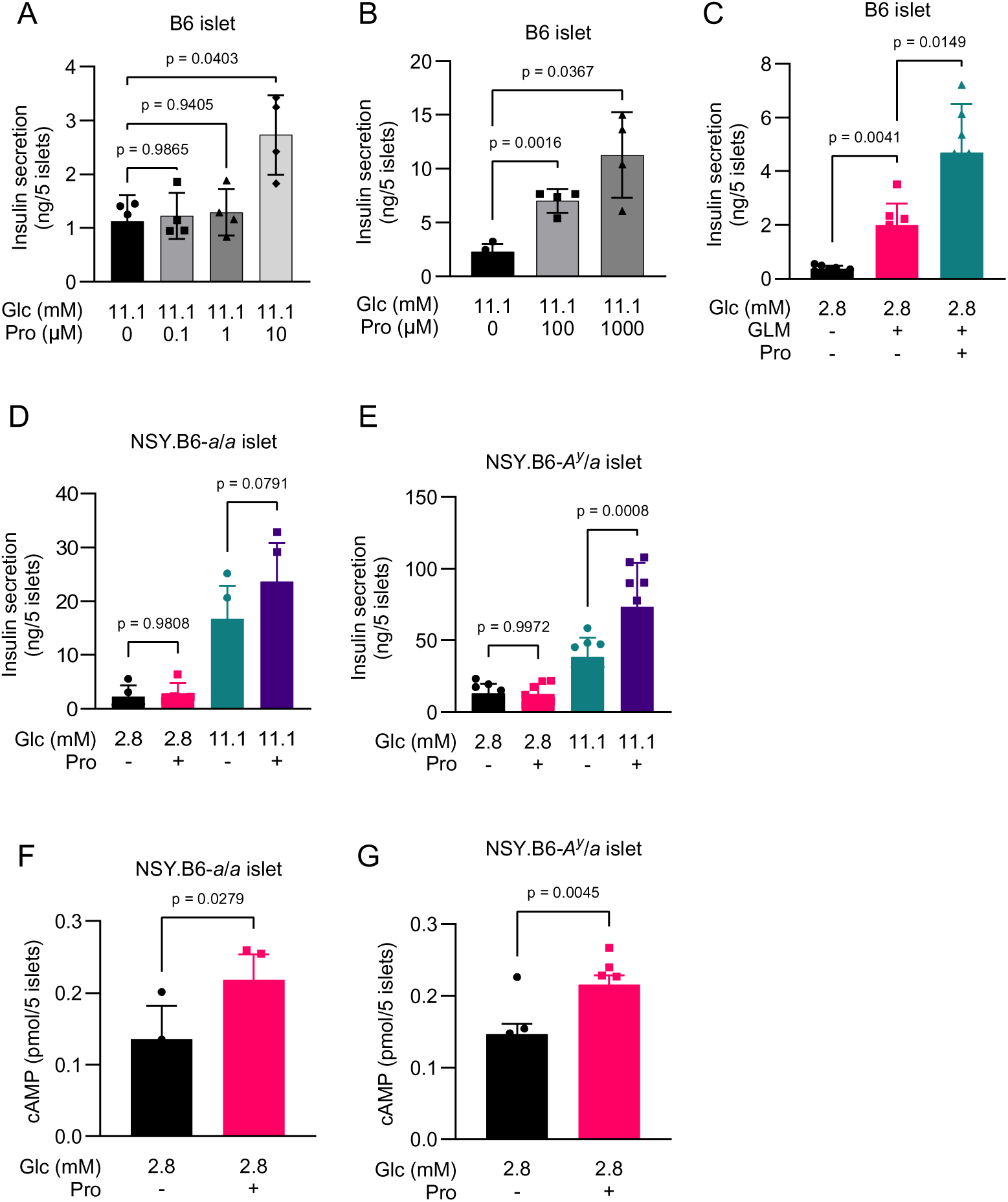
β-Blocker-stimulated cAMP elevation and insulin secretion in isolated mouse islets. A. –B Dose-dependent effect of propranolol on glucose-induced insulin secretion in B6 mouse islets. One mouse was used for each experiment. n = 4. C. Effect of propranolol on glimepiride-induced insulin secretion in B6 mouse islets. One mouse was used. n = 7. D. –E. Effect of propranolol on insulin secretion in islets isolated from (D) NSY.B6*-a*/*a* mice (n = 5) and (E) NSY.B6*-A^y^*/*a* mice (n = 8). Islets were pooled from two mice for each genotype. F. –G. Effect of propranolol on cAMP levels in islets isolated from (F) NSY.B6*-a*/*a* mice (n = 4–5) and (G) NSY.B6-Ay/a mice (n = 7) were pooled from two mice. Islets were pooled from two mice for each genotype. Glc, glucose. Propranolol (Pro): 100 μM. Glimepiride (GLM): 1 μM. Statistical comparisons were performed using Welch’s one-way ANOVA with Dunnett’s post-hoc test for A– E, and Welch’s unpaired two-tailed t-test for F and G.

### Propranolol potentiates Ca^2+^ response via PKA/RYR2 pathway

To clarify whether propranolol-SIS involves potentiation of intracellular Ca^2+^ ([Ca^2+^]_i_), we conducted Fluo-4 imaging in MIN6-K8 cells. Propranolol enhanced the glimepiride-induced increase in [Ca^2+^]_i_ (Figure 4, A–C). Glimepiride alone elicited oscillatory [Ca^2+^]_i_ waves, whereas a combination of glimepiride and propranolol induced a rapid increase in [Ca^2+^]_i_, followed by rapid oscillations (Figure 4B).

**Figure 4.**
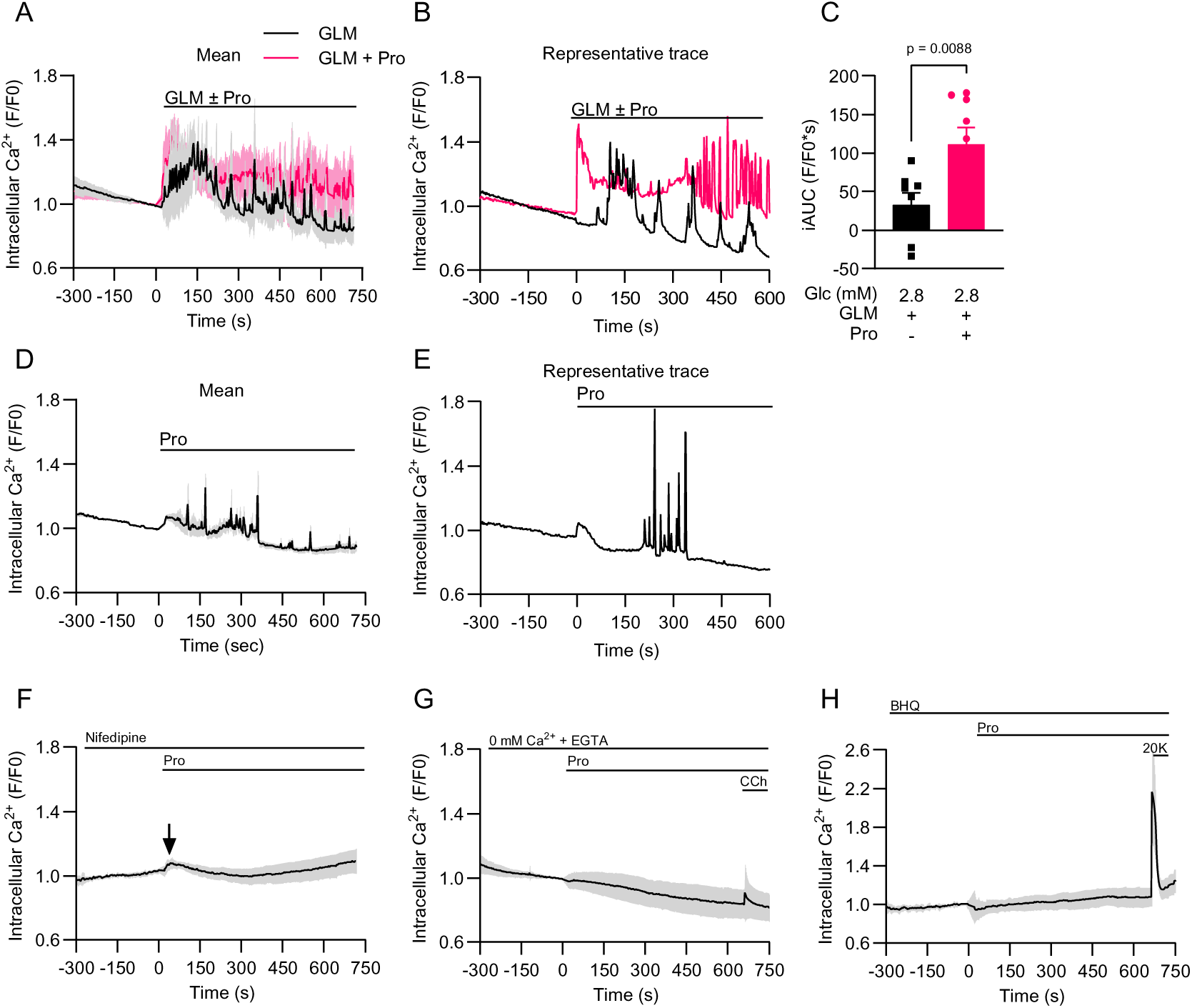
Characterization of propranolol-stimulated Ca^2+^ response in MIN6-K8 cells. Intracellular Ca^2+^ levels were measured using Fluo-4. All experiments were performed using MIN6-K8 cells in the presence of 2.8 mM glucose. The averaged time courses of normalized fluorescence intensity (F/F0) are indicated in A, D, and F–H. Representative traces are indicated in B and E. A. –C. Effects of propranolol on the glimepiride-induced Ca^2+^ response. GLM: n = 8; GLM + Pro: n = 9. C, The magnitude of Ca^2+^ responses was quantified as the incremental area under the curve (iAUC). D. –E. Propranolol-stimulated Ca^2+^ response. n = 7. F. Effect of nifefdipine (100 μM) on the glimepiride-induced Ca^2+^ response. n = 12. G. Propranolol-stimulated Ca^2+^ response in extracellular Ca^2+^-free conditions. n = 17. EGTA: 100 μM. Carbachol (CCh, 50 μM) was added as a positive control at the end of the recording. The black arrow indicates the [Ca^2+^]_i_ increase at 0 min. H. Effect of BHQ (15 μM) on glimepiride-induced Ca^2+^ response. n = 19. 20 mM K^+^ (20K) was added as a positive control at the end of the recording. All experiments were conducted using MIN6-K8 cells in the presence of 2.8 mM glucose. Data are presented as mean ± SD. Glc, glucose. The reagents were added to achieve the following final concentrations unless otherwise specified: glimepiride (GLM), 1 μM; and propranolol (Pro), 100 μM. Statistical comparisons were made using Welch’s unpaired two-tailed t-test for C.

Notably, propranolol alone elicited a transient [Ca^2+^]_i_ increase at 0 min, followed by sporadic spikes (Figure 4, D–E). This propranolol-stimulated Ca^2+^ response was further investigated to elucidate the involvement of extracellular Ca^2+^ and intracellular Ca^2+^ stores. Nifedipine, an inhibitor of L-type voltage-dependent Ca^2+^ channels (VDCC), suppressed the sporadic [Ca^2+^]_i_ spikes but not the [Ca^2+^]_i_ increase at 0 min indicated by the black arrow (Figure 4F), suggesting that the [Ca^2+^]_i_ spikes represent Ca^2+^ influx through L-type VDCCs. In contrast, removal of extracellular Ca^2+^ from the stimulation buffer inhibited both the [Ca^2+^]_i_ increase at 0 min and subsequent [Ca^2+^]_i_ spikes (Figure 4G). Depletion of intracellular Ca^2+^ stores with 2,5-di-tert-butylhydroquinone (BHQ), an inhibitor of sarcoplasmic/endoplasmic reticulum Ca^2+^ ATPase (SERCA), exerted the same effect (Figure 4H). These results suggest that the initial [Ca^2+^]_i_ increase involves both [Ca^2+^]_i_ mobilization from the endoplasmic reticulum (ER) and extracellular Ca^2+^ influx and is required for the later VDCC-mediated [Ca^2+^]_i_ spikes. Importantly, nifedipine, removal of extracellular Ca^2+^, and BHQ also abolished propranolol-SIS (Figure S3, A–C), demonstrating that the propranolol-stimulated Ca^2+^ response is essential for propranolol-SIS.

The mobilization of Ca^2+^ from the ER is mediated by specific ion channels, such as ryanodine receptors (RYRs) and IP_3_ receptors (IP_3_Rs), which release Ca^2+^ into the cytosol in response to signaling events (Guerrero-Hernández et al., 2020). The involvement of RYRs was investigated using dantrolene, an RYR inhibitor. Dantrolene negated the capacity of propranolol to potentiate the glimepiride-induced Ca^2+^ response in MIN6-K8 cells (Figure 5, A–B). Dantrolene paradoxically increased glimepiride-induced insulin secretion, consistent with a previous report (Figure S3D) (Pian-Smith et al., 1986); however, it decreased propranolol-IIS, as expressed by the fold change (Figure 5C), suggesting that the propranolol-stimulated Ca^2+^ response and insulin secretion are dependent on RYR. It has been reported that the open probability of RYR2 is increased by intracellular cAMP through PKA-mediated RYR2 phosphorylation at Ser-2808 (Marx et al., 2000; Wehrens et al., 2006). Remarkably, phosphorylation of RYR2 at Ser-2808 was substantially induced by propranolol alone and further enhanced by the addition of glimepiride (Figure 5, D–E). These findings suggest that the propranolol-stimulated Ca^2+^ response is mediated by PKA-induced RYR2 phosphorylation, corroborating the observation that propranolol-SIS is dependent on PKA activity (Figure 2F).

**Figure 5.**
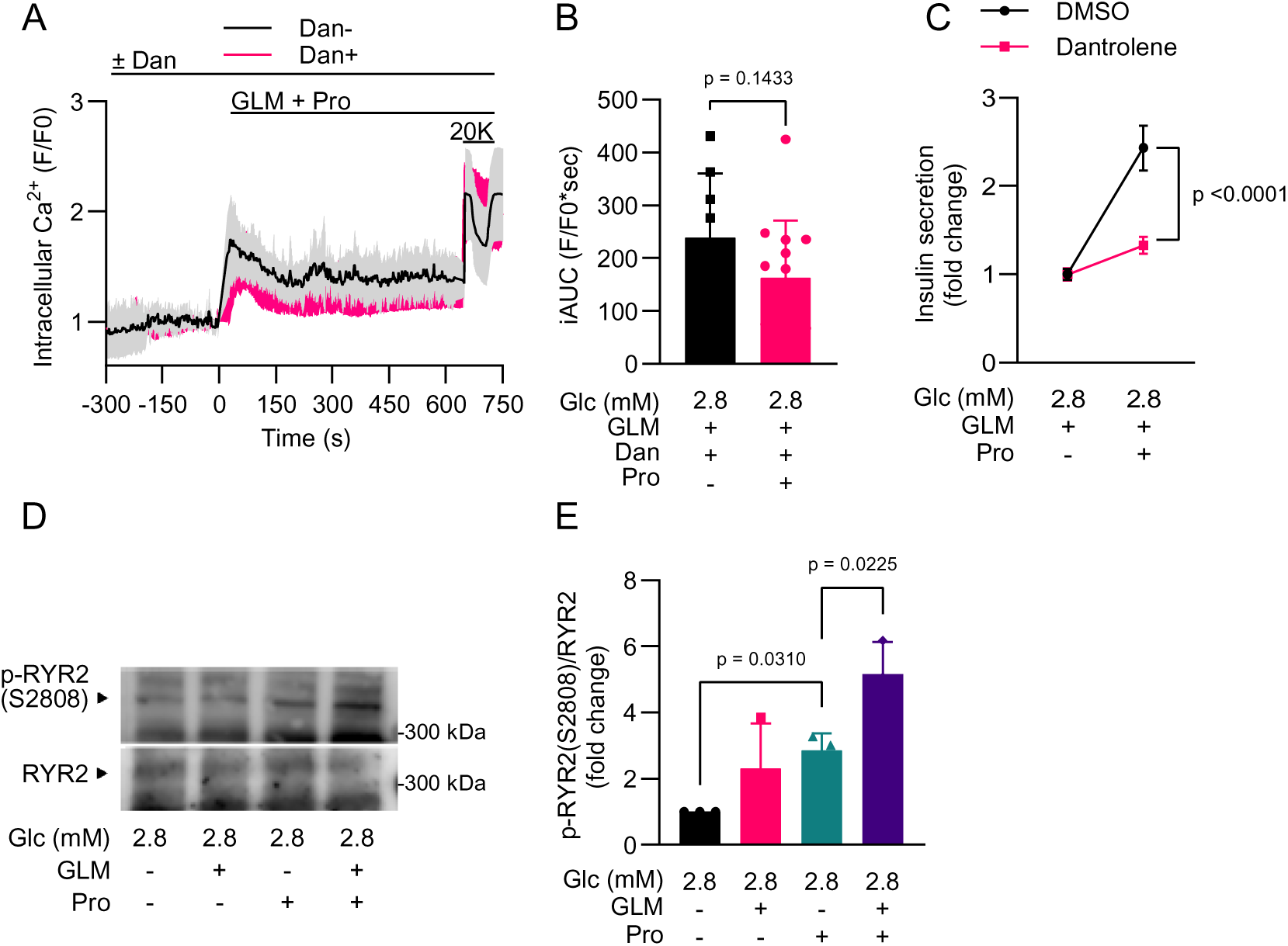
Involvement of ryanodine receptor in propranolol-stimulated Ca^2+^ response in MIN6-K8 cells. A. –B. Effect of dantrolene on the propranolol-stimulated Ca^2+^ response in the presence of glimepiride. The averaged time courses of the normalized fluorescence intensity of Fluo-4 (F/F0) are indicated. Dan-: n = 9; Dan+: n = 14. GLM + Pro: n = 9. C, 20 mM K^+^ (20K) was added as a positive control at the end of the recording. The magnitude of Ca^2+^ responses was quantified as the incremental area under the curve (iAUC) between 0 and 600 s. C. Effect of dantrolene on propranolol-SIS. The data are presented as fold change. See Figure S3D for the original values. D. –E. The effects of glimepiride and propranolol on RYR2 phosphorylation at Ser-2808 were assessed by immunoblotting. The intensity of phosphorylated RYR2 (Ser-2808) was normalized to total RYR2 (Ser-2808) and expressed as fold change over 2.8 mM glucose in E. n = 3. All experiments were performed using MIN6-K8 cells in the presence of 2.8 mM glucose. Data are presented as mean ± SD. The reagents were added to achieve the following final concentrations, unless otherwise specified: dantrolene (Dan), 100 μM; glimepiride (GLM), 1 μM; and propranolol (Pro), 100 μM. Statistical comparison was made using Welch’s unpaired two-tailed t-test for B and C, and Welch’s one-way ANOVA with Dunnett’s post-hoc test for E.

The involvement of IP_3_Rs was investigated using xestospongin C, an IP_3_R inhibitor (Gafni et al., 1997), and siRNA-mediated knockdown of IP_3_ receptor 1 (Itpr1). Xestospongin C marginally decreased propranolol-SIS (Figure S3, E–F). However, depletion of *Itpr1* by approximately 50% at the transcript level did not affect propranolol-SIS, as expressed by the fold-change (Figure S3, G–I). These findings suggest that IP_3_ receptors play a limited, if any, role in propranolol-SIS.

### Involvement of TRPM5 in propranolol-stimulated Ca^2+^ response

We investigated the mechanism by which PKA-mediated [Ca^2+^]_i_ mobilization from the ER leads to VDCC activation. Interestingly, a reduction in extracellular Na^+^ from 150 mM to 30 mM suppressed the propranolol-stimulated Ca^2+^ response and insulin secretion in MIN6-K8 cells (Figure 6, A–B), suggesting the involvement of Na^+^-permeable channels. Transient receptor potential cation channel subfamily M member 4 (TRPM4) and member 5 (TRPM5) are Ca^2+^-activated channels that permeate monovalent cations, including Na^+^ and K^+^. They are gated by [Ca^2+^]_i_, most notably Ca^2+^ from the ER (Zhang et al., 2007; Gonzales et al., 2010). Notably, TRPM5 has been implicated in the augmentation of glucose-induced depolarizing currents in β-cells (Colsoul et al., 2010), suggesting its potential role as a mediator linking [Ca^2+^]_i_ mobilization from the ER to depolarization. To elucidate the involvement of TRPM4 and TRPM5, we conducted siRNA-mediated knockdown, which depleted their mRNA levels by approximately 50% (Figure S4, A–B). Knockdown of *Trpm4* and *Trpm5* led to slight and moderate decreases in propranolol-SIS, respectively (Figure 6C, Figure S4C). Furthermore, triphenylphosphine oxide (TPPO), a TRPM5 inhibitor (Palmer et al., 2010), inhibited the later Ca^2+^ spikes and insulin secretion (Figure 6, D–E). These findings suggest that TRPM5 is involved in the later phase of the propranolol-stimulated Ca^2+^ response, contributing to propranolol-SIS.

**Figure 6.**
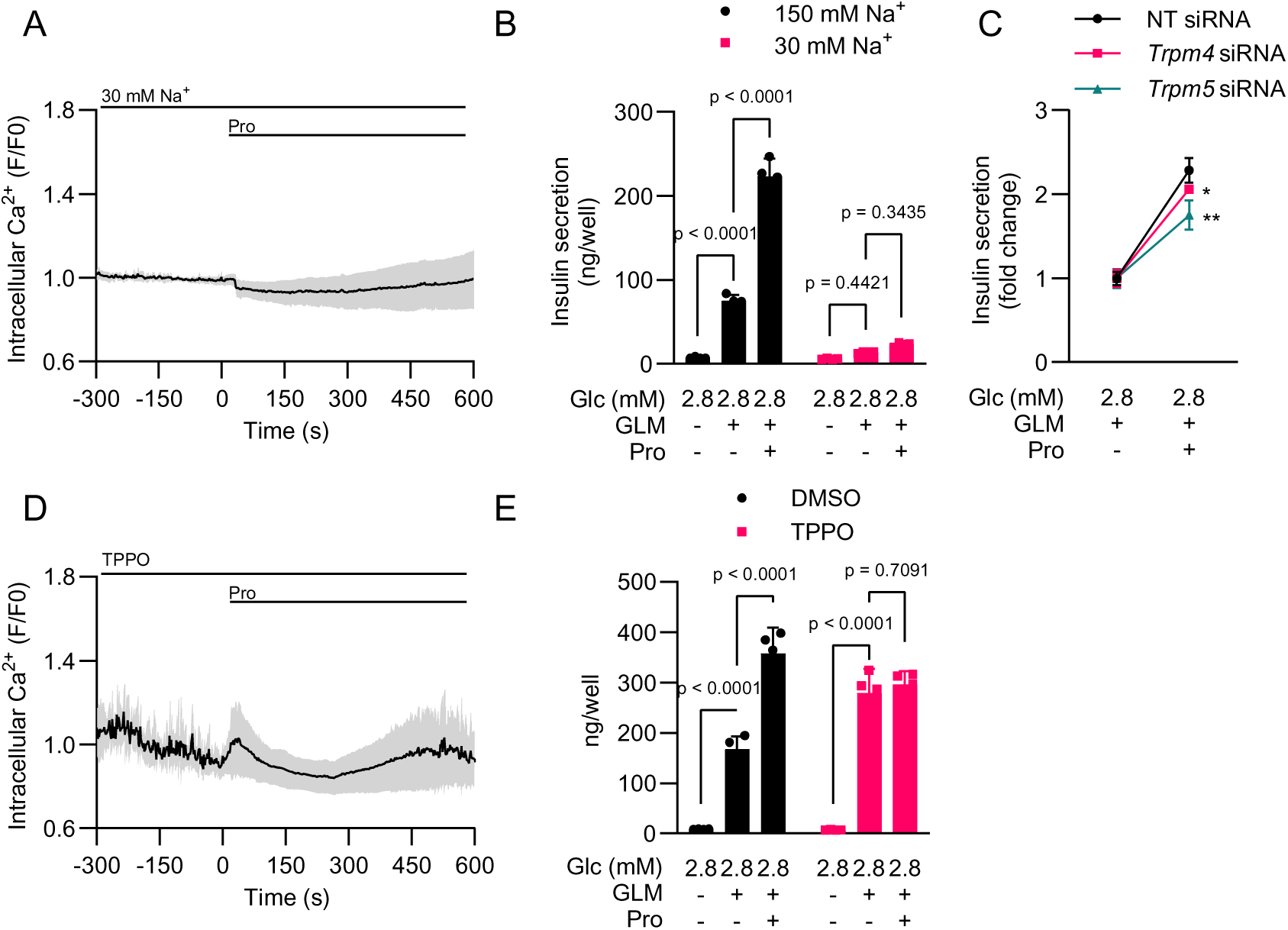
Involvement of TRPM4 and TRPM5 in propranolol-stimulated Ca^2+^ response in MIN6-K8 cells. A. Effect of low extracellular sodium concentration (30 mM) on the propranolol-stimulated Ca^2+^ response. The averaged time course of the normalized fluorescence intensity of Fluo-4 (F/F0) is indicated. n = 12. B. Effect of low extracellular sodium concentration (30 mM) on propranolol-SIS. n = 4. C. Effects of *Trpm4* and *Trpm5* knockdown on propranolol-SIS. n = 4. The data are presented as fold change. See Figure S4C for the original values. *p < 0.05, **p < 0.01 versus NT (non-targeting) siRNA. D. Effect of triphenylphosphine oxide on propranolol-stimulated Ca^2+^ response. The averaged time course of the normalized fluorescence intensity of Fluo-4 (F/F0) is indicated. n = 12. E. Effect of triphenylphosphine oxide on propranolol-SIS. n = 4. All experiments were performed using MIN6-K8 cells in the presence of 2.8 mM glucose. Data are presented as mean ± SD. The reagents were added at the following final concentrations unless otherwise specified: glimepiride (GLM), 1 μM; propranolol (Pro), 100 μM; and triphenylphosphine oxide (TPPO), 100 μM. Statistical comparisons were made using Welch’s one-way ANOVA with Dunnett’s post-hoc test for C and two-way ANOVA with Šídák’s post-hoc test for B and E.

### **α**2-adrenoceptors are involved in cAMP elevation by propranolol

Using previously published in-house RNA-seq datasets (Hashim et al., 2018), we investigated the expression of adrenoceptor subtypes in β-cells. MIN6-K8 cells exclusively expressed α2A-adrenoceptors, while the other subtypes were undetectable (Table 1). Similar patterns were observed in B6 mouse islets, which exhibited high expression of α2A-adrenoceptors and minimal levels of α1, β1, and β2 receptors, potentially originating from non-β-cells. These findings align with previous research indicating that mouse β-cells exclusively express α2A-adrenoceptors (Rorsman and Ashcroft, 2018).

**Table 1.**
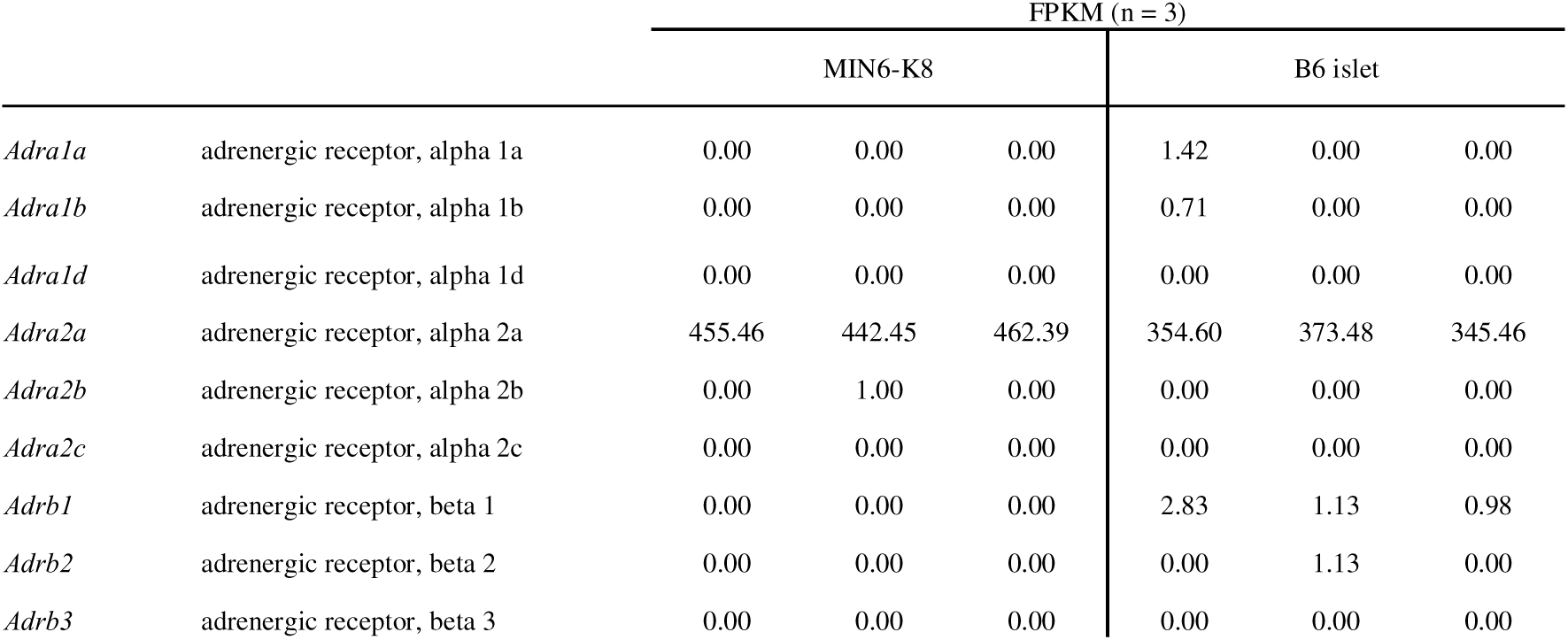
Expression profiles of adrenoceptors in MIN6-K8 cells and B6 mouse islets. Source data: DDBJ Sequence Read Archive (DRA) accession number DRA006332 (Hashim et al., 2018). n = 3 for each sample type.

Interestingly, Pretreatment with atipamezole (α2-adrenoceptor antagonist) and phentolamine (α1/2-adrenoceptor antagonist) attenuated propranolol-stimulated cAMP elevation, as expressed by the fold-change (Figure 7, A–B). These inhibitors also abolished propranolol-SIS (Figure 7, C–D). Although phentolamine elevated glimepiride-induced insulin secretion, presumably due to its direct inhibitory effect on K_ATP_ channels (Plant and Henquin, 1990; Proks and Ashcroft, 1997), it potently suppressed propranolol-SIS (Figure 7D). To further validate the involvement of α2A-adrenoceptors, we performed siRNA- mediated Adra2a knockdown. Although the knockdown efficiency was only moderate at the protein level (28%, as shown in Figure S5, A–B), propranolol-SIS was decreased as expressed by the fold change (Figure 7E, S5C). These results indicate that α2-adrenoceptors are the only adrenoceptor subtype expressed in mouse β-cells, and they play a permissive role in cAMP elevation and insulin secretion by β-blockers.

**Figure 7.**
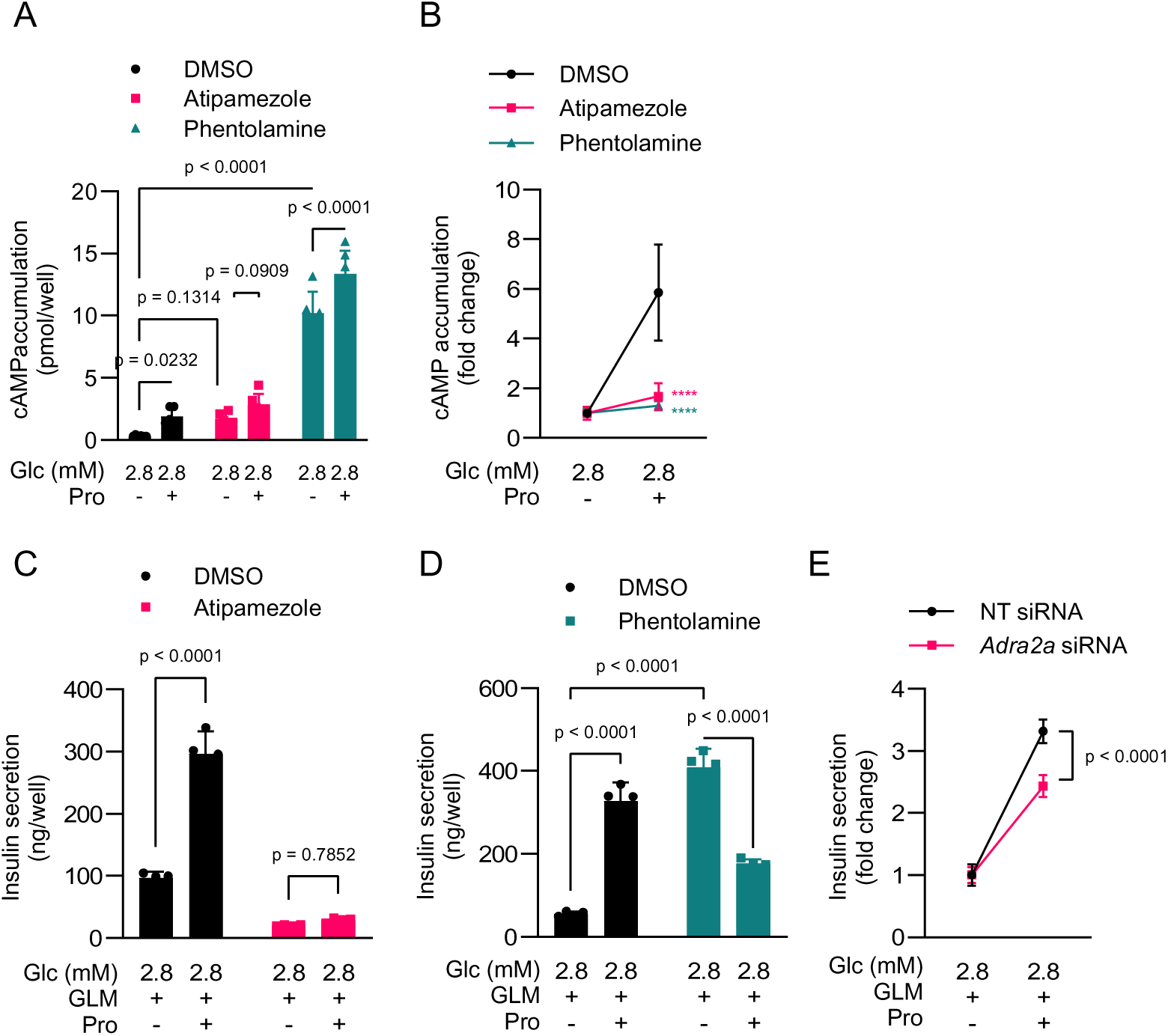
Involvement of α2-adrenoceptors in propranolol-stimulated cAMP elevation and insulin secretion. A. –B. Effects of α2-adrenoceptor inhibitors on intracellular cAMP accumulation. n = 6. The data **are** presented **as (A)** original value and as (B) fold change over 2.8 mM glucose alone. C. –D. Effects of α2-adrenoceptor inhibitors, (C) atipamezole and (D) phentolamine, on propranolol-SIS. n = 4. were used. n = 4. E. Effect of α2A-adrenoceptor knockdown on propranolol-SIS. n = 4. The data are presented as fold change. See Figure S5C for the original values. All experiments were performed using MIN6-K8 cells in the presence of 2.8 mM glucose. Data are presented as mean ± SD. Glc, glucose. The reagents were added to achieve the following final concentrations, unless otherwise specified: glimepiride (GLM), 1 μM; propranolol (Pro), 100 μM; phentolamine, 100 μM; and atipamezole, 250 μM. Statistical comparisons were made using Welch’s one-way ANOVA with Dunnett’s post-hoc test for B (****p < 0.0001 versus DMSO), two-way ANOVA with Šídák’s post-hoc test for A and C, two-way ANOVA with uncorrected Fisher’s LSD for D, and Welch’s unpaired two-tailed t- test for E.

## Discussion

This study reports a paradoxical off-target effect of β-blockers on mouse pancreatic β-cells: β-blockers increased cAMP levels, [Ca^2+^]_i_, and insulin secretion despite the absence of β- adrenoceptors in these cells. We used propranolol to elucidate the signaling pathway of β- blocker-stimulated insulin secretion, which involves cAMP/PKA/RYR2/TRPM5.

Despite the widespread use of β-blockers in clinical and experimental settings, their concentrations vary significantly. Clinical pharmacokinetic research indicates that in healthy subjects, the maximum plasma concentration (Cmax) of propranolol rarely surpasses 1 μM after a single oral administration (Abdu-Aguye et al., 1986; Taegtmeyer et al., 2014; Kim et al., 2022). The therapeutic range for propranolol is typically considered to be 192 to 385 nM (0.05 to 0.10 μg/mL), with toxic effects occurring above 7 μM (2 μg/mL) and fatal outcomes at levels exceeding 11 μM (3 μg/mL) (Kerns, 2007; Sharifpour et al., 2022). Similarly, the in vivo Cmax of bisoprolol generally remains below 100 nM (Li et al., 2012; Tjandrawinata et al., 2012).

In experimental contexts, propranolol exhibits near-complete inhibition of β2- adrenoceptors at nanomolar levels (IC50 < 1 nM) (Boursier et al., 2020). Nevertheless, some cell experiments have used propranolol at concentrations exceeding 100 μM. These studies involved various cell types, including hemangioma cells (Munabi et al., 2015; Kum and Khan, 2014), clear cell renal cell carcinoma cells (Shepard et al., 2018), and PC3 and MCF7 cancer cell lines (Reyes-Corral et al., 2019). Our findings suggest that caution should be exercised when using high concentrations of β-blockers, as they may produce off-target effects, contrary to conventional β-blocker pharmacology. Indeed, Reyes-Corral et al. reported that 50 μM propranolol triggered CICR in PC3 and MCF7 cancer cell lines, although they did not examine cAMP levels (Reyes-Corral et al., 2019).

Signaling pathways linking cAMP and insulin secretion have been studied extensively (Seino and Shibasaki, 2005; Tengholm and Gylfe, 2017). However, this is the first report of the PKA/RYR2/TRPM5 pathway. Notably, propranolol evoked a Ca^2+^ response in MIN6-K8 cells even under low-glucose conditions, in which cells are electrically silent. The Ca^2+^ response comprised an immediate transient increase at 0 min and subsequent spikes, the latter indicative of Ca^2+^ influx through L-type VDCCs. We demonstrated that the initial increase in [Ca^2+^]_i_ involves [Ca^2+^]_i_ mobilization through RYR2, which is facilitated by PKA-mediated RYR2 phosphorylation at Ser 2808. Supporting our findings, it has been suggested that RYR2 activation in β-cells requires cAMP-dependent phosphorylation (Islam et al., 1998). However, that report also indicates that cAMP-dependent phosphorylation alone is insufficient to mobilize Ca^2+^ from the ER when cytosolic [Ca^2+^] is low (Islam et al., 1998; Islam, 2002); instead, it only sensitizes RYR2 to Ca^2+^-induced Ca^2+^ release (CICR) by releasing the channel from Mg^2+^-mediated inhibition (Hain et al., 1994) or FKBP12.6 (FK-506-binding protein 12.6) binding (Marx et al., 2000). Thus, it is reasonable to postulate that propranolol-stimulated cAMP triggers both initial Ca^2+^ influx and RYR2 phosphorylation, leading to CICR, reflected in the transient rise in [Ca^2+^]_i_ at 0 min. The mediator of propranolol-stimulated initial Ca^2+^ influx is currently unclear. However, given that it was inhibited by depletion of extracellular Ca^2+^ and Na^+^, but not by nifedipine, cAMP-activated nifedipine-insensitive Ca^2+^ channels, such as T-type VDCCs, may be involved (Louiset et al., 2017; Sharma et al., 2023). A similar mechanism has been proposed for glucagon-like peptide 1 (GLP-1)-stimulated insulin secretion: GLP-1-stimulated cAMP production leading to CICR through a dual effect on Ca^2+^ influx and RYR2 sensitization (Holz et al., 1999).

TRPM5 and/or TRPM4 have been implicated in background inward currents produced by GLP-1 (Shigeto et al., 2015) and fructose (Kyriazis et al., 2012). TRPM5 functions downstream of Gαq- or gustducin-coupled GPCR signaling cascades, wherein PLCβ activation triggers Ca^2+^ release from the IP_3_ receptor and protein kinase C (PKC) activation, both of which facilitate TRPM5 gating (Shigeto et al., 2015). Our findings suggest that TRPM5 can also be activated downstream of the cAMP/PKA signaling cascade, highlighting its versatile role in linking GPCR signaling and membrane depolarization. Small background inward currents, including those from TRPM5, are incapable of inducing depolarization unless the majority of β-cell KATP channels are closed (Rorsman and Ashcroft, 2018). This mechanism may explain why membrane depolarization plays a permissive role in propranolol-SIS.

The exact mechanism by which β-blockers elevate cAMP levels remains unclear. In our cellular model, α2-adrenoceptors were the sole adrenoceptor subtype expressed and played a permissive role in propranolol-induced cAMP elevation and insulin secretion. This finding indicates that propranolol may potentially interact with α2-adrenoceptors, although such an effect has not been previously documented. Notably, isoproterenol, a β-adrenoceptor agonist, was found to inhibit GIIS at high concentrations (10–50 μM), and this effect was prevented by the preinhibition of α2-adrenoceptors (Zielmann et al., 1985).

In summary, we revealed that β-blockers paradoxically increase cAMP levels in β-cells, highlighting the variable nature of β-blocker effects, which are potentially influenced by concentration and cell type. These findings underscore the importance of careful consideration in experimental design when working with β-blockers.

## Abbreviations

AC: adenylyl cyclase
BHQ: 2,5-di-*tert*-butylhydroquinone
cAMP: adenosine 3’,5’-Cyclic Monophosphate
CICR: Ca^2+^-induced Ca^2+^ release
ER: endoplasmic reticulum
FKBP: FK-506-binding protein
GPCR: G-protein coupled receptor
GLP-1: glucagon-like peptide
GSIS: glucose-stimulated insulin secretion
iAUC: incremental area under the curve
IP1: inositol 1-phosphate
IP_3_: inositol 1,4,5-trisphosphate
K_ATP_ channel: ATP-sensitive K^+^ channel
Propranolol-SIS: propranolol-stimulated insulin secretion
PKA: protein kinase A
PKC: protein kinase C
PLCβ: phospholipase Cβ
SERCA: sarcoplasmic/endoplasmic reticulum Ca^2+^ ATPase
TRPM: Transient receptor potential cation channel subfamily M
VDCC: voltage-dependent Ca^2+^ channel
β-blocker-SIS: β-blocker-stimulated insulin secretion

## Acknowledgements

The authors extend their gratitude to President Yutaka Seino of Kansai Electric Power Hospital for his generous support in this research. The authors thank Yoshikazu Hoshino of Hoshino Laboratory Animals, Inc. for providing NSY.B6*-A^y^* mice. The authors thank Shihomi Sakai, Asami Yamaguchi, and Megumi Akiyama for their excellent technical assistance.

## Data Availability

The authors declare that all data supporting the findings of this study are contained within the paper and Supplemental Data.

## Authorship Contributions

*Participated in research design*: Murao.

*Conducted experiments*: Murao, Morikawa, Shimomura, and Maejima.

*Performed data analysis*: Murao.

*Wrote or contributed to the writing of the manuscript*: Murao, Seino, Yamada, and Suzuki.

## Footnotes

This study was supported by JSPS KAKENHI Grant Numbers JP22K20869 and JP23K15401 for N.M. Research grants for N.M. were provided by the Japan Association for Diabetes Education and Care, Daiwa Securities Foundation, Suzuken Memorial Foundation, Japan Diabetes Foundation, The Hori Sciences and Arts Foundation, Manpei Suzuki Diabetes Foundation, and Fujita Academy.

---

**Figure S1.**
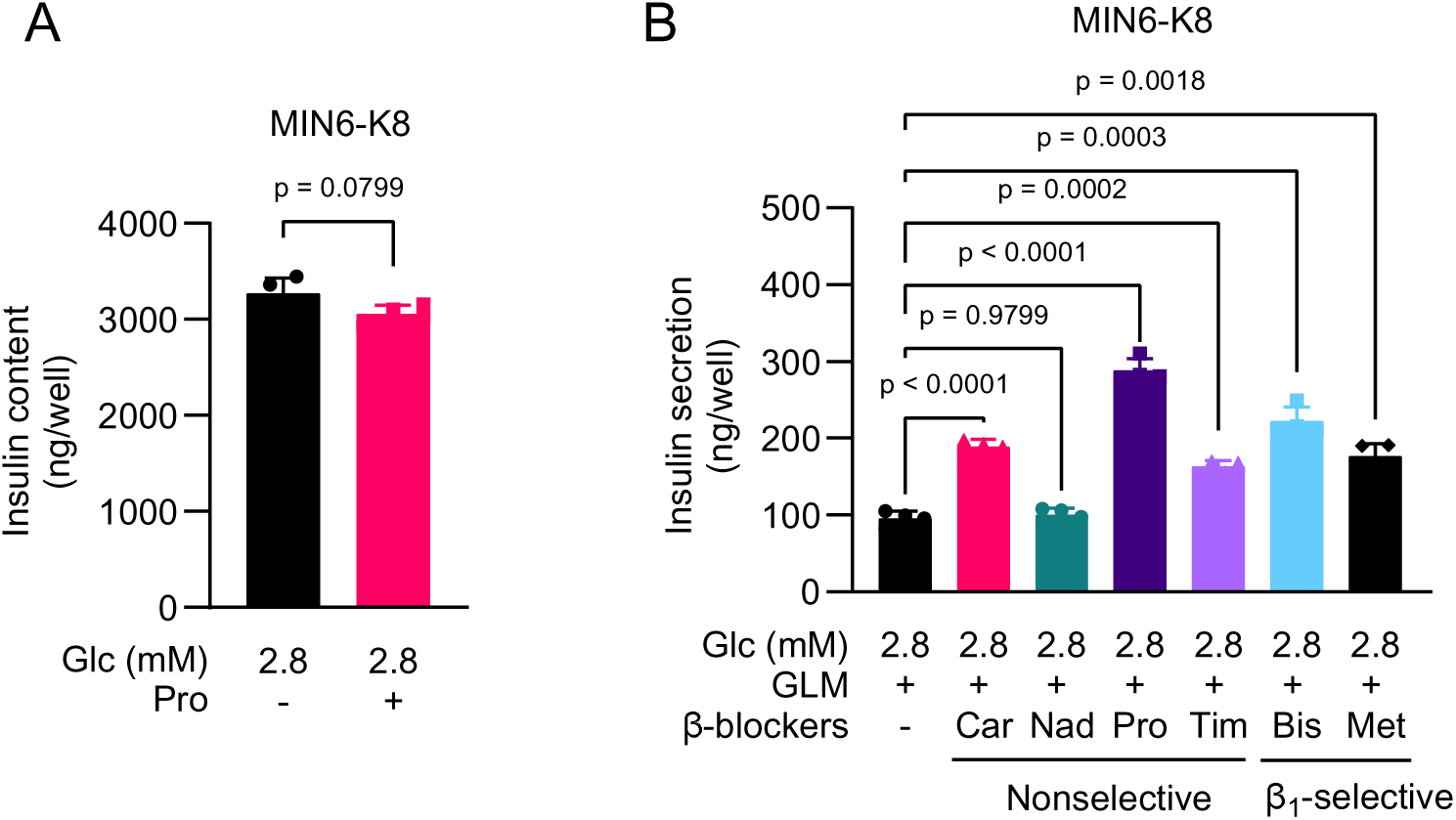
Effect of β-blockers on insulin secretion and content in MIN6-K8 cells. A. Effect of propranolol on intracellular insulin content. n = 4. B. Effect of various non-selective and β1-selective β-blockers on glimepiride-induced insulin secretion in MIN6-K8 cells. Car, carteolol; Nad, nadolol; Pro, propranolol; Tim, timolol; Bis, bisoprolol; Met, metoprolol. All β-blockers were used at a concentration of 100 μM. n = 4. All experiments were performed using MIN6-K8 cells in the presence of 2.8 mM glucose. Glimepiride (GLM): 1 μM. Propranolol (Pro): 100 μM. Bisoprolol (Bis): 100 μM. Data are presented as mean ± SD. Statistical comparisons were made using Welch’s one-way ANOVA with Dunnett’s post-hoc test for C, and Welch’s unpaired two-tailed t-test for A, and two-way ANOVA with Šídák’s post-hoc test for B.

**Figure S2.**
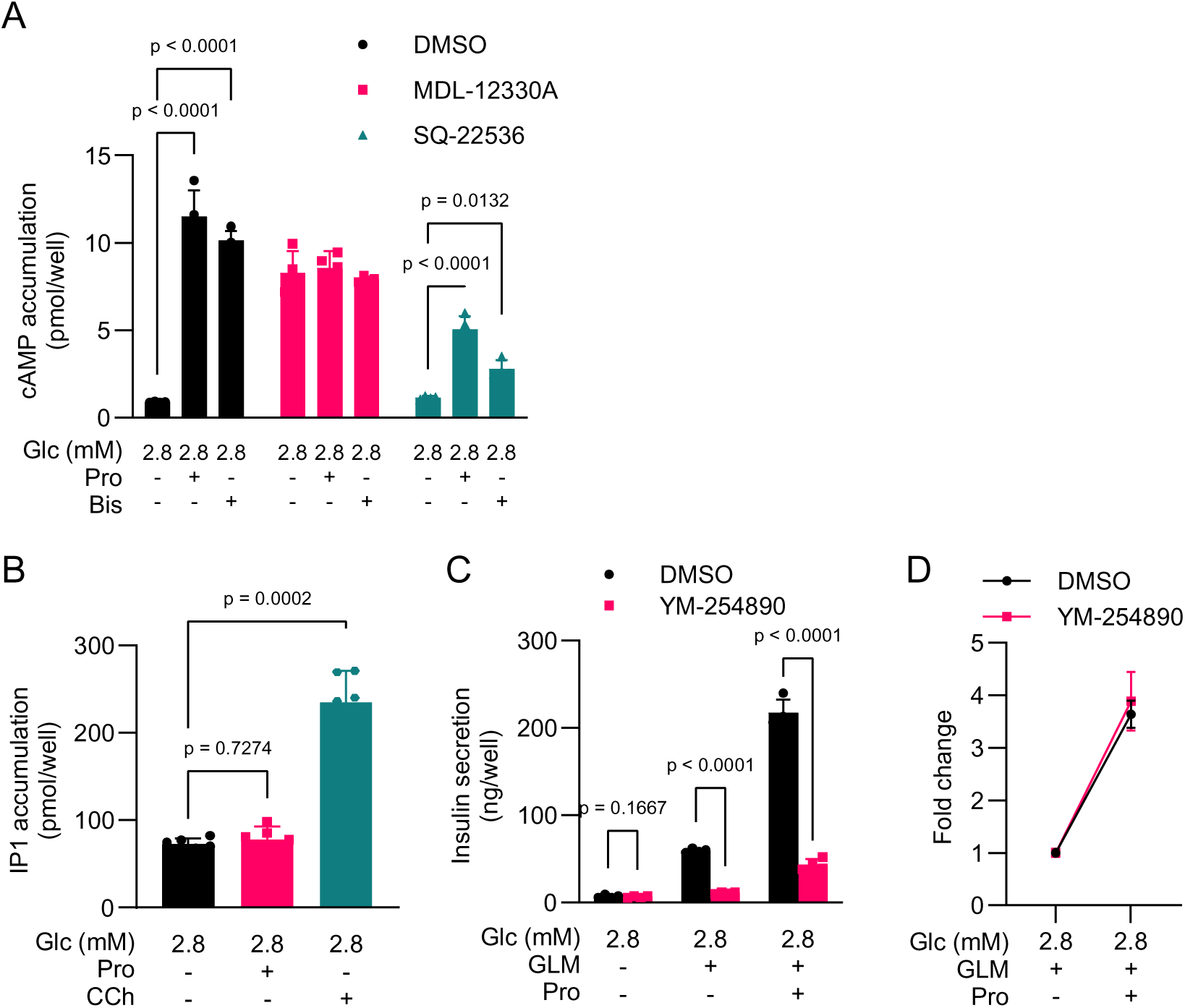
Involvement of G protein signaling pathways in propranolol-stimulated insulin secretion in MIN6-K8 cells. A. Effects of AC inhibitors on β-blocker-stimulated cAMP production. n = 4., MDL-12330A – 10 μM, SQ-22536 – 1 mM. B. Effect of propranolol on intracellular IP1 accumulation. n = 6. Carbachol (CCh, 50 μM) was used as the positive control. C. –D. Effect of YM-254890 (100 nM) on propranolol-SIS. n = 4. The data are presented in their original value in C and as fold change over glimepiride in D. All experiments were performed using MIN6-K8 cells. Glc, glucose. Glimepiride (GLM): 1 μM. Propranolol (Pro): 100 μM. Bisoprolol (Bis): 100 μM. Data are presented as the mean ± SD. Statistical comparisons were made using two-way ANOVA with Šídák’s post-hoc test for A, Welch’s one-way ANOVA with Dunnett’s post-hoc test for B, and Welch’s unpaired two-tailed t-test for C.

**Figure S3.**
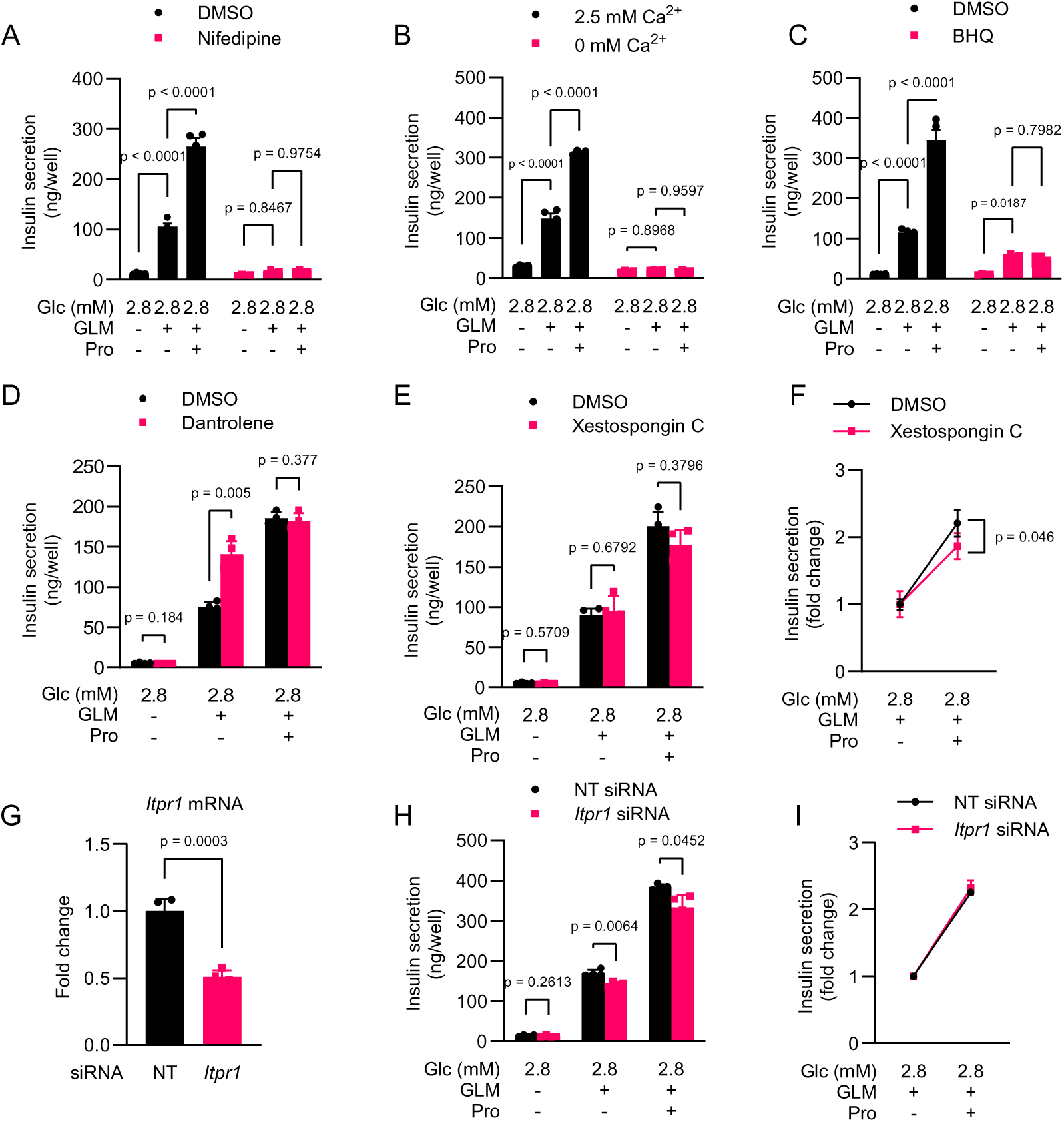
Involvement of intracellular and extracellular Ca^2+^ in propranolol-stimulated insulin secretion in MIN6-K8 cells. A. Effect of nifedipine (100 μM) on propranolol-SIS. n = 4. B. Propranolol-SIS under extracellular Ca^2+^-free conditions. n = 4. EGTA (100 μM) was added to the Ca^2+^-free buffer. C. Effect of BHQ (15 μM) on propranolol-SIS. n = 4. D. Effect of dantrolene (100 μM) on propranolol-SIS. n = 4. E. –F. Effect of xestospongin C (2 μM) on propranolol-SIS. n = 4. The data are presented in their original value in E and as fold change over glimepiride in F. G. The knockdown efficiency of *Itpr1* was assessed using RT-qPCR. The mRNA levels were normalized to those in siNT (non-targeting siRNA)-treated cells. n = 4. H. –I. Effect of *Itpr1* knockdown on propranolol-SIS. n = 4. The data are presented in their original value in H and as fold change over glimepiride in I. All experiments were performed using MIN6-K8 cells. Glc, glucose. Glimepiride (GLM): 1 μM. Propranolol (Pro): 100 μM. Data are presented as mean ± SD. Statistical comparisons were made using two-way ANOVA with Šídák’s post-hoc test for A–C and Welch’s unpaired two-tailed t-test for D–H.

**Figure S4.**
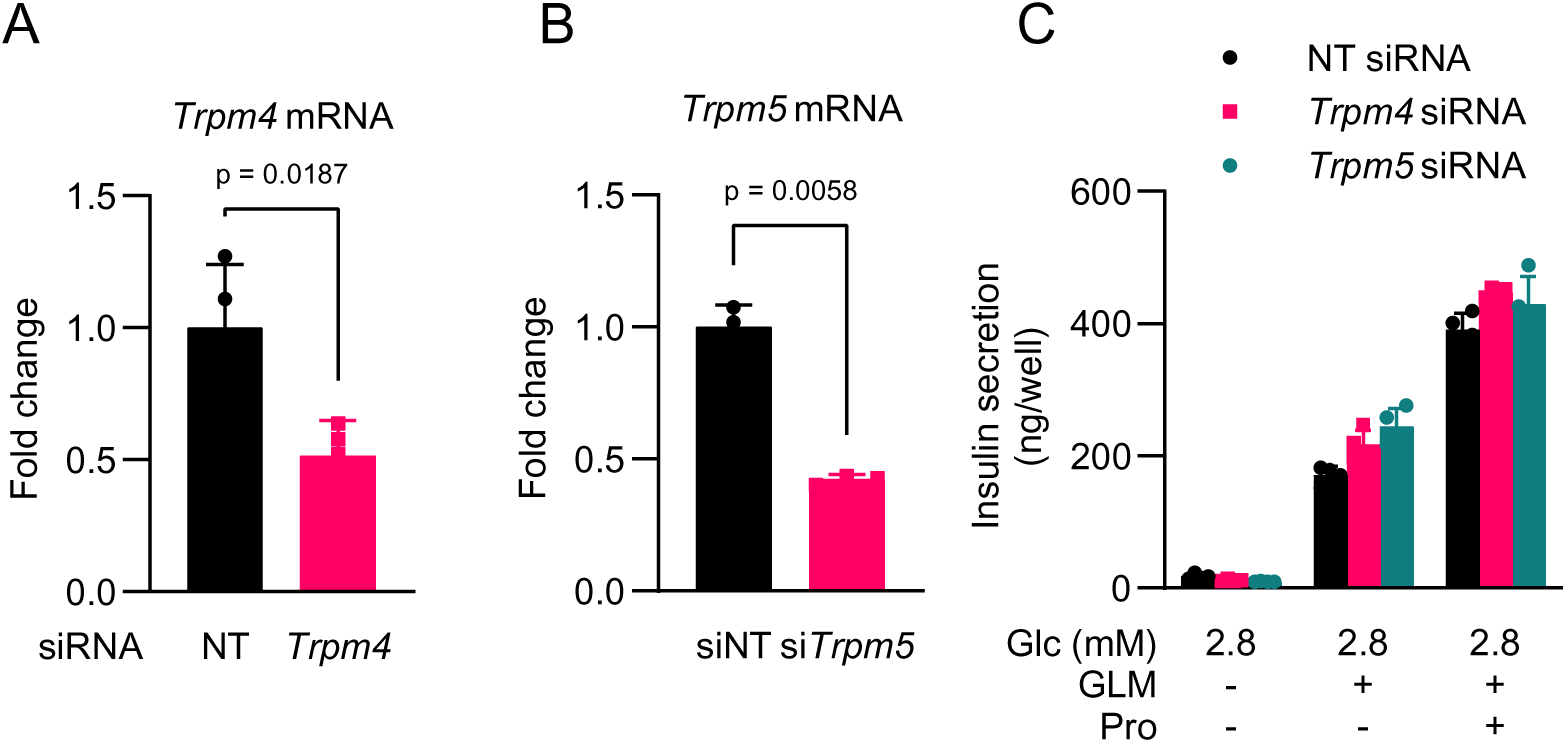
Effects of *Trpm4* and *Trpm5* knockdown on propranolol-stimulated insulin secretion in MIN6-K8 cells. A. –B. The knockdown efficiencies of *Trpm4* and *Trpm5* were assessed using RT-qPCR. The mRNA levels were normalized to those in siNT (non-targeting siRNA)-treated cells. n = 4. C. Effect of *Trpm4* and *Trpm5* knockdown on propranolol-SIS. n = 4. All experiments were performed using MIN6-K8 cells. Glc, glucose. Glimepiride (GLM): 1 μM. Propranolol (Pro): 100 μM. Data are presented as mean ± SD. Statistical comparisons were made using Welch’s unpaired two-tailed t-test.

**Figure S5.**
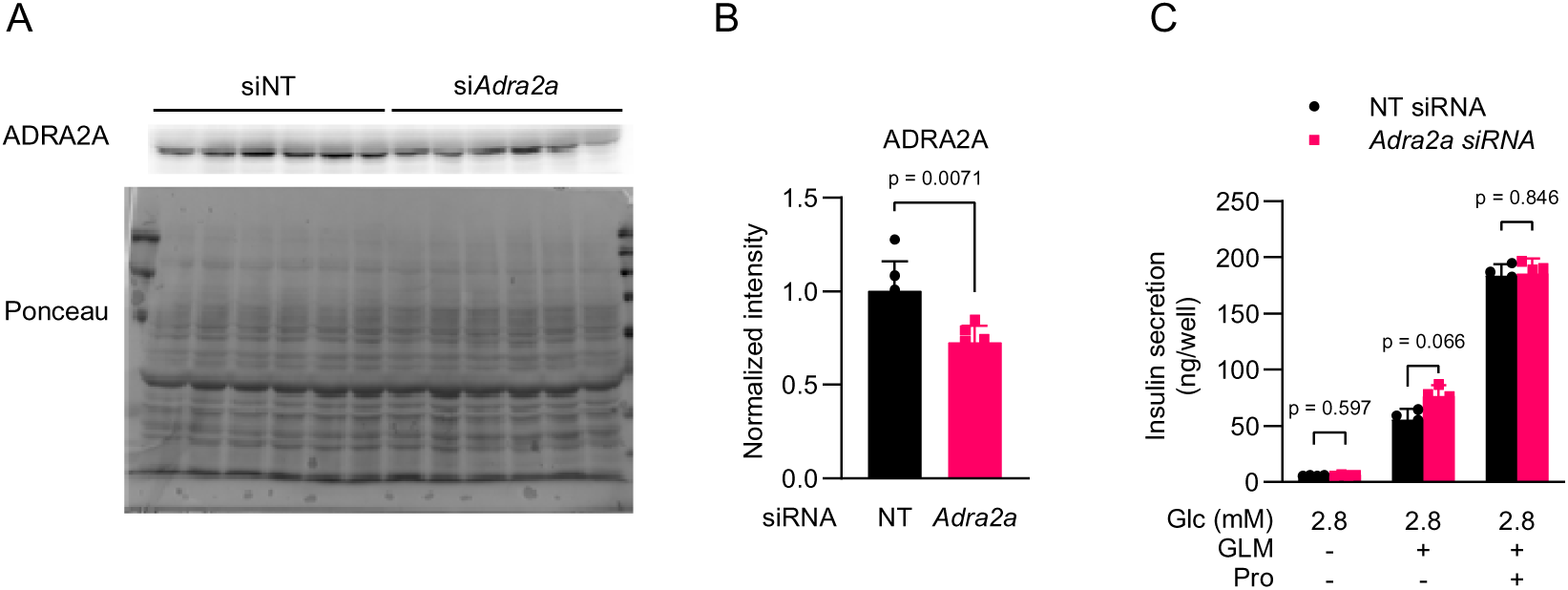
Effects of *Adra2a* knockdown on propranolol-stimulated insulin secretion in MIN6-K8 cells. A. –B. Knockdown efficiency of *Adra2a* was assessed by immunoblotting. (B) The intensity of ADRA2A was normalized to that of total protein visualized by Ponceau-S staining and expressed as fold change over 2.8 mM glucose. n = 6. C. Effect of *Adra2a* knockdown on propranolol-SIS. n = 4. All experiments were performed using MIN6-K8 cells. Glc, glucose. Glimepiride (GLM): 1 μM. Propranolol (Pro): 100 μM. Data are presented as the mean ± SD. Statistical comparisons were made using Welch’s unpaired two-tailed t-test.

